# Microbiota-derived short-chain fatty acids mediate *Candida albicans* gastrointestinal colonization resistance

**DOI:** 10.1101/2025.09.08.674977

**Authors:** Animesh A. Mishra, Laura A. Coughlin, Nicole Poulides, Jiwoong Kim, Xiaowei Zhan, Shuheng Gan, Sebastian E. Winter, Christina M. Zarek, Lora V. Hooper, Andrew Y. Koh

## Abstract

The gut microbiota plays a critical role in constraining *Candida albicans* (*Ca*) colonization of the gastrointestinal (GI) tract, a key precursor to disseminated fungal infection in immunocompromised hosts. Depletion of commensal microbiota increases *Ca* burden and promotes dissemination, yet the mechanisms of microbiota-mediated *Ca* colonization resistance remain poorly defined. Here, we show that gut microbiota-derived short-chain fatty acids (SCFAs) directly inhibit *Ca* growth by impairing hexose uptake, disrupting central carbon metabolism, and inducing intracellular acidification. *In vivo*, SCFAs enhance *Ca* colonization resistance only in the presence of an intact gut microbiome, which is required to drive SCFA-induced taxonomic shifts that further augment resistance. Commensal microbiota lacking SCFA production exhibit diminished capacity to restrict *Ca* colonization, while prebiotic therapy that increases luminal SCFA levels enhances *Ca* clearance. These findings define a critical microbiota–metabolite mechanism underlying *Ca* colonization resistance and suggest strategies to modulate GI fungal burden and prevent invasive disease.

## Introduction

*Candida albicans* (*Ca*), a leading human fungal pathogen, can exist as a pathobiont in the gastrointestinal (GI) tract and/or genitourinary tract^1^. In healthy individuals, asymptomatic *Ca* carriage is the norm. In immunocompromised hosts, however, *Ca* can cause severe disseminated and fatal infections^2,3^.

Specific bacterial taxa promote colonization resistance against *Ca* in the mammalian GI tract^4,5^. For example, commensal anaerobes *Bacteroides thetaiotamicron* and *Blautia producta* promote colonization resistance against *Ca* in the murine GI tract, a finding corroborated in human stem cell transplant patients colonized with *Candida*^4^. These commensals induce intestinal epithelial expression of the transcription factor, HIF-1*α*, which leads to production of the *Ca*-active cathelicidin antimicrobial peptide, LL-37/CRAMP^4^. Yet, a multitude of other mechanisms likely contribute to gut microbiota-driven *Ca* colonization resistance.

One common feature of both *B. theta* and *B. producta* is their ability to produce short-chain fatty acids (SCFAs)^6^. SCFAs – including acetate, propionate, and butyrate – are key bacterial-derived metabolites with broad immunomodulatory and antimicrobial properties. SCFAs are notable for promoting immune tolerance^7^, inducing antimicrobial peptide production from intestinal epithelial cells^8^, and directly inhibiting bacterial growth, particularly among Enterobacteriaceae^9,10^.

Recent studies demonstrate that butyrate shapes intestinal oxygen dynamics by promoting colonocyte oxygen consumption, thereby maintaining epithelial hypoxia and restricting the growth of facultative anaerobes (including *Ca*) that proliferate under aerobic conditions in the gut^11,12^. However, the direct effects of SCFAs on *Ca* growth, metabolism, and GI colonization remain poorly understood.

Here, we show that SCFAs directly inhibit *Ca* growth *in vitro* by impairing hexose uptake, disrupting central carbon metabolism, and inducing intracellular acidification. *In vivo*, SCFA-mediated *Ca* colonization resistance is contingent upon the presence of gut bacteria. Commensal microbiota unable to produce SCFAs exhibit diminished capacity to reduce *Ca* colonization. Moreover, prebiotic therapy that enhances gut SCFA levels promotes *Ca* clearance. These findings establish SCFAs as critical mediators of microbiota-driven *Ca* colonization resistance.

## Results

### Short-chain fatty acids (SCFA) inhibit *Candida albicans* (*Ca*) growth *in vitro*

The most commonly studied *Ca* strain SC5314 cannot sustainably colonize the GI tract of mice with intact gut microbiomes^13,14^. Thus, preclinical models often utilize antibiotics to deplete endogenous gut bacteria^13,15^. Antibiotics that deplete anaerobic species, many of which produce SCFA, are most effective in promoting *Ca* colonization^5^. Indeed, mice treated with anti-anaerobic antibiotics (penicillin/streptomycin ^5^) for 7 days had significantly higher *Ca* GI colonization levels (**Fig 1a)**, consistent with our prior reports^5,15-17^ and had significantly lower levels of the SCFAs acetic acid (AA), butyric acid (BA), and propionic acid (PA) in cecal fecal contents (**Fig 1b**).

**Figure 1.**
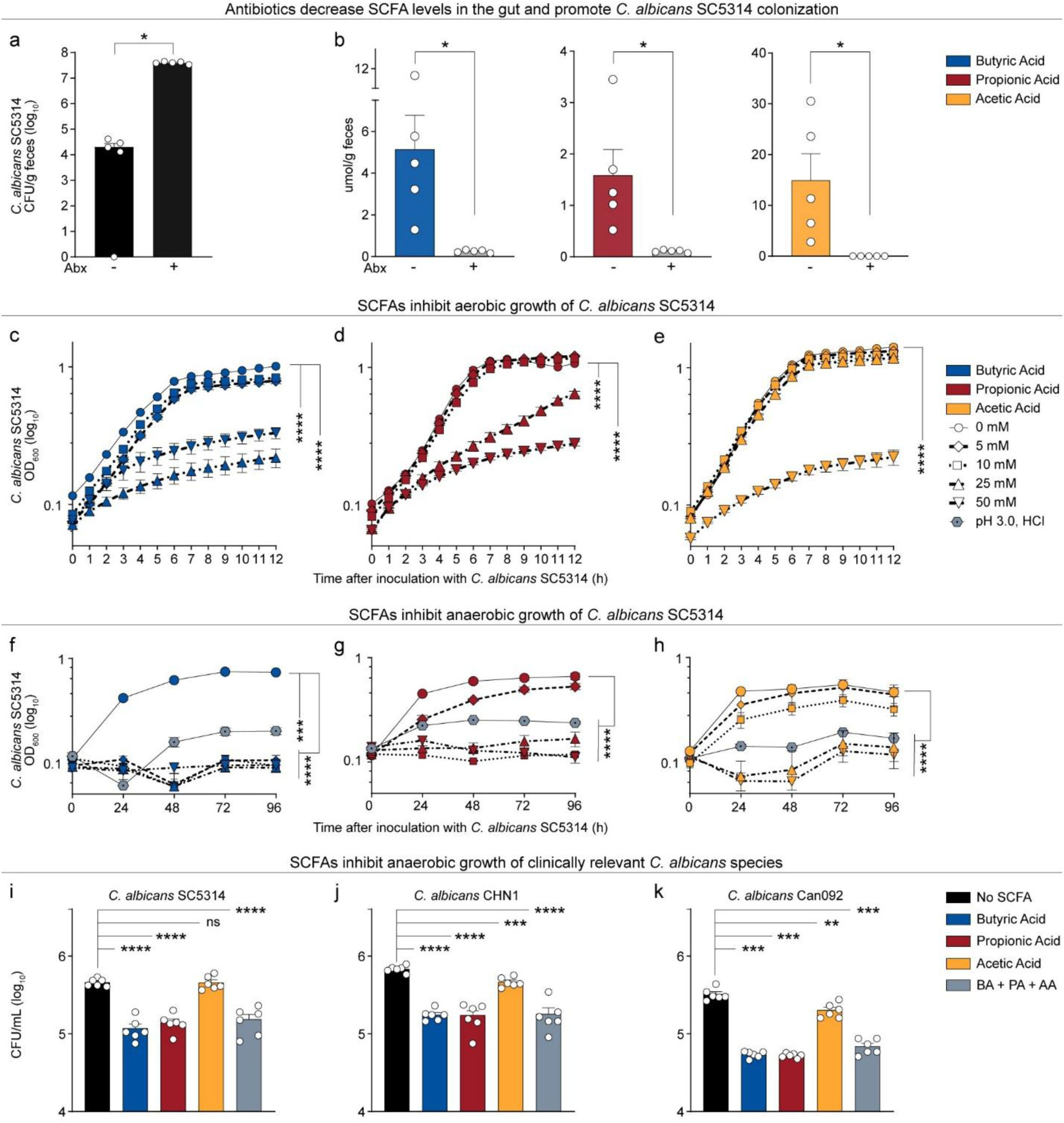
Short-chain fatty acids (SCFA) inhibit *Candida albicans* (*Ca*) growth *in vitro*. **(a)** *Ca* SC5314 colonization levels in C57BL/6 (female, 6-8 weeks, Jackson) mice ± antibiotics ad libitum in water for one week and then orally gavaged with 2×10^8^ CFU of *Ca* SC5314, as determined by cultured enumeration of fecal homogenates. Fecal pellets collected 14 days post inoculation. Fecal homogenates serially diluted and plated on YPD plates with gentamicin and vancomycin for *Ca* CFU enumeration. **(b)** SCFA levels in cecal contents (as measured by GC-MS) from mice (female, 6-8 weeks, Jackson) treated ± antibiotics *ad libitum* in water. (**a-b**) Abx, antibiotics (penicillin/streptomycin). Points = results from individual animals. n=5 per group. Bars = mean ± SEM. Statistical analysis by (**a**) Mann-Whitney and (**b**) unpaired t-test. \**P* < 0.05. *Ca* SC5314 aerobic **(c-e)** and anaerobic **(f-h)** growth curves ± SCFA treatment. *Ca* cultured for 12 hours in YPD aerobically or anaerobically at 37°C with varying concentrations of butyric acid (BA), propionic acid (PA), and acetic acid (AA). No treatment control was YPD adjusted to pH=3.0 with HCl. OD_600_ measured every hour for aerobic studies and every 24 h for anaerobic studies. Bars = mean ± SEM from three independent experiments. Linear mixed effect model with Kruskal-Wallis test. \*\*\**P* < 0.001, \*\*\*\**P* < 0.0001. Enumeration of live *C. albicans* clinical isolates — **(i)** SC5314, **(j)** CHN1, and **(k)** Can092 — grown in YPD ± SCFA for 4 hours under anaerobic conditions. <u>S</u>erial dilution of cultures on YPD plates with vancomycin and gentamycin. Bars = mean ± SEM CFU recovered from cultures from two independent experiments. Each experiment had three technical replicates per condition. Unpaired t-test. \*\**P* < 0.01, \*\*\**P* < 0.001, \*\*\*\**P* < 0.0001, ns, not significant.

We then investigated the effect of SCFA on *Ca* growth *in vitro*. *Ca* SC5314 aerobic growth was significantly reduced with increasing concentrations of BA, PA, or AA in the media (**Fig. 1c, 1d, 1e)**. BA and PA inhibited *Ca* growth most significantly at concentrations ≥25 mM (**Fig. 1c, 1d**) whereas AA required concentrations of 50 mM for greatest growth inhibition (**Fig. 1e**). Since SCFA acidifies media, we tested if the growth inhibition was attributable to pH reduction alone. *Ca* aerobic growth was not impeded by changes in pH alone (YPD titrated with HCl, pH range of 3-6) (**Fig S1**), consistent with observations that *Ca* can robustly modulate extracellular pH^18^.

Oxygen concentrations are scarce or absent in the gastrointestinal tract, particularly the distal gut^11,19^. We assessed the effect of SCFA on *Ca* growth under anaerobic conditions and tested the effect of acidification alone (pH 3.0, HCl). All three SCFAs were able to inhibit the growth of SC5314 under anaerobic conditions more effectively than under aerobic conditions (**Fig. 1f, g, h**). In anaerobic conditions, acidification of the media alone inhibited *Ca* growth, but not to the extent of higher SCFA concentrations. Finally, antibiotic treatment not only lowers SCFA levels in the distal gut but also increases oxygen concentrations at/near intestinal mucosal surfaces^11,20,21^. This combined reduction in SCFAs and rise in oxygen^12^ may promote *Ca* colonization, as the fungus grows more efficiently under aerobic conditions, even in the presence of SCFAs.

We also quantified SCFA growth inhibition by enumeration of cultured *Ca* grown in presence or absence of SCFA under anaerobic conditions. The number of viable *Ca* was significantly lower when grown in the presence of BA and PA, but not when grown with AA (**Fig. 1i**). Interestingly, all three SCFAs (mixed in proportions physiologically relevant to the mammalian GI tract; 13 mM AA, 5 mM PA, 7 mM BA^7^) were able to inhibit the growth of SC5314 as effectively as BA and PA alone (**Fig. 1i**).

Strain-specific differences can significantly impact *Ca*’s ability to colonize the murine GI tract. In contrast to *Ca* SC5314, some clinical *Ca* strains (CHN1^17^) are able to sustainably colonize the murine GI tract without the use of antibiotics^17^. As observed with *Ca* SC5314, all SCFAs were effective in inhibiting clinical strains Can092^5^ and CHN1 growth under anaerobic conditions, with BA and PA more effective than AA (**Fig. 1j, k**). These data suggest that SCFA-induced *Ca* growth inhibition is not a strain-specific phenomenon.

### SCFAs remodel the *C. albicans* transcriptome and suppress metabolic and growth pathways

Transcriptional profiling (*Ca* SC5314, *in vitro* aerobically, with or without 50mM AA, BA, or PA) revealed extensive remodeling of the *Ca* transcriptome in response to BA and PA, with ∼562–600 genes downregulated and ∼845–865 genes upregulated, of which >87-94% were shared between BA and PA, respectively (**Supplemental Tables, RNAseq tabs**). In contrast, AA elicited minimal transcriptional responses, with only 5 genes upregulated and 194 downregulated, most of them unique to AA (**Fig. 2a, b**). Gene pathway enrichment analysis of the BA/PA-shared differentially expressed genes (GO processes, **Fig. 2a, b**; GO functions, **Fig. S2a, b**; **Supplemental Tables, GO tabs**) revealed coordinated suppression of translational processes (Pol I/III, rRNA/tRNA processing) and ribosome biogenesis, and reprogramming of metabolism toward survival pathways. Concurrently, an upregulation of glycolysis, trehalose/glycogen reserves, redox enzymes, and nutrient salvage/transport was observed, consistent with a shift toward an energy-conserving, stress-tolerant state under SCFA-induced growth inhibition.

**Figure 2.**
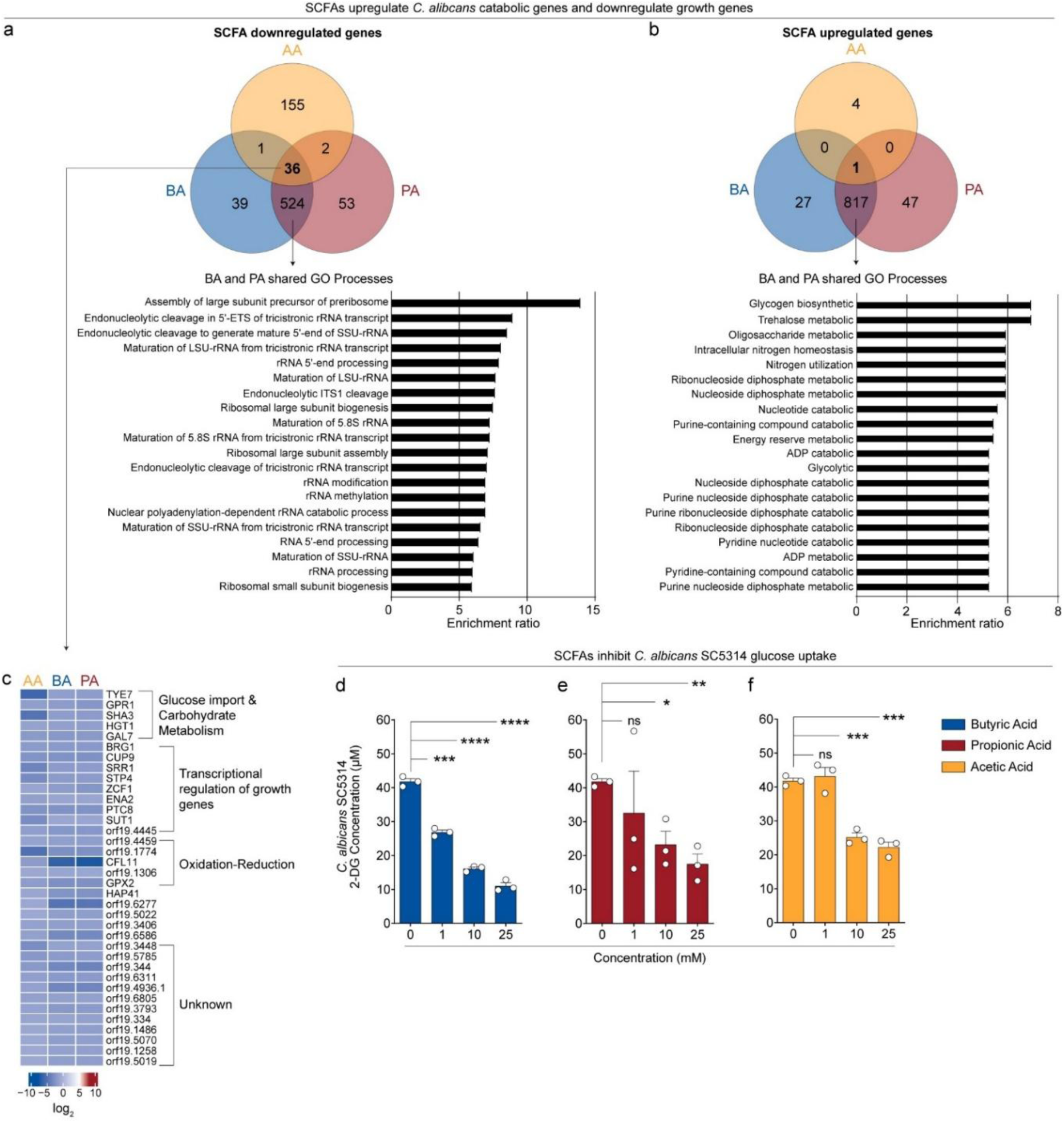
SCFAs inhibit hexose uptake and disrupt central carbon metabolism in *Ca*. **(a–c)** Transcriptomic profiling of SC5314 cells cultured ±50 mM butyrate (BA), propionate (PA), or acetate (AA) for 30 min under aerobic conditions. Total RNA was extracted from three biological replicates per condition and analyzed by RNA-seq. Differentially expressed genes were defined as >2-fold change versus untreated control with adjusted *p* < 0.05 (false discovery rate correction). **(a)** Number of unique and shared downregulated genes and pathway enrichment analysis (Gene Ontology, top 20 biological processes; Bonferroni-adjusted *p* < 0.05) of BA/PA-shared downregulated genes. **(b)** Number of unique and shared upregulated genes and pathway enrichment analysis (Gene Ontology, top 20 biological processes; Bonferroni-adjusted *p* < 0.05) of BA/PA-shared upregulated genes. **(c)** Heatmap and KEGG pathway enrichment for genes downregulated across all SCFA treatments. (**d–f**) 2-deoxyglucose (2-DG) uptake assay in *Ca* SC5314 cells exposed to (**d**) BA, (**e**) PA or (**f)** AA treatment. Intracellular 2-DG was quantified fluorometrically (Ex/Em = 537/585 nm) using a standard curve. Bars = mean ± SEM from three independent experiments. Unpaired *t*-test. *P* < 0.05, \**P* < 0.01, \*\**P* < 0.001, \*\*\**P* < 0.0001, ns, not significant.

We examined the 36 genes significantly downregulated by all three SCFAs (>2-fold decreased, adjusted *p* < 0.05), consistent with the observation that each SCFA inhibits *Ca* growth *in vitro*. These included key regulators of nutrient (glucose) sensing (*GPR1^22,23^*), hexose uptake (*SHA3^24,25^*, *HGT1^26-28^*), carbohydrate metabolism (*TYE7*, *GAL7*^29-33^), oxidation-reduction (*CFL11^34^, GPX2^35,36^*), and growth and morphology factors (*BRG1^37^*, *CUP9^38,39^*, *SRR1^40^*, *STP4^41^*, *ZCF1^42,43^*, *ENA2^44^*, *PTC8^45^*, *SUT1^46,47^*) (**Fig. 2a, c**). Of note, *Ca* mutants lacking *TYE7* fail to colonize the murine GI tract^31^. While dispensable for basal growth^34-36^, redox genes may become critical under stress conditions, such as SCFA exposure where redox homeostasis is disrupted. Like the BA/PA gene pathway enrichment analyses, these data suggest that SCFA exposure suppresses a regulatory axis extending from higher-order transcriptional control (*BRG1*, *STP4*, *CUP9*) through membrane transporters (*HGT1*), ultimately converging on core metabolic and growth pathways. This coordinated suppression may drive a starvation-like state that contributes to SCFA-mediated growth inhibition.

### SCFAs impair glucose uptake in *C. albicans*

To test this hypothesis, we used 2-deoxyglucose (2-DG), a non-metabolizable glucose analog that is taken up by cells and thus serves as a functional readout for glucose uptake capacity^48,49^. SCFA treatment caused a dose-dependent decrease in intracellular 2-DG levels (**Fig. 2d–f)**. BA exerted the strongest inhibition, at SCFA concentrations as low as 1 mM (**Fig. 2d**), followed by PA (**Fig. 2e**). In contrast, AA had the weakest effect (**Fig. 2f**). These findings corroborate the transcriptome data and identify impaired glucose uptake as a mechanism by which SCFAs inhibit *Ca* growth.

### SCFAs disrupt central carbon and amino acid metabolism in *C. albicans*

To further define how SCFAs affect *Ca* metabolism, we profiled intracellular metabolites of SCFA-treated *Ca* SC5314 cells using hydrophilic interaction liquid chromatography spectrometry-mass spectrometry (HILIC-MS), an unbiased platform that detects 591 metabolites. BA and PA induced strikingly similar metabolic signatures, while AA diverged for ∼50% of detected metabolites (**Fig 3a-b**; **Supplemental Tables, “Metabolomics_diff abundant” tab**). Among shared changes, α-ketoglutarate, a tricarboxylic acid cycle (TCA) NADH-producing intermediate, was depleted under all SCFA treatments, while adenosine 2’,3’-cyclic phosphate accumulated, peaking with BA and PA. This metabolite, a positional isomer of cAMP^50,51^, has been linked to mitochondrial dysfunction^50,51^ and implicated in stress responses in *E.coli*^52^, plants^53^, and mammalian cells^51,54^.

**Figure 3.**
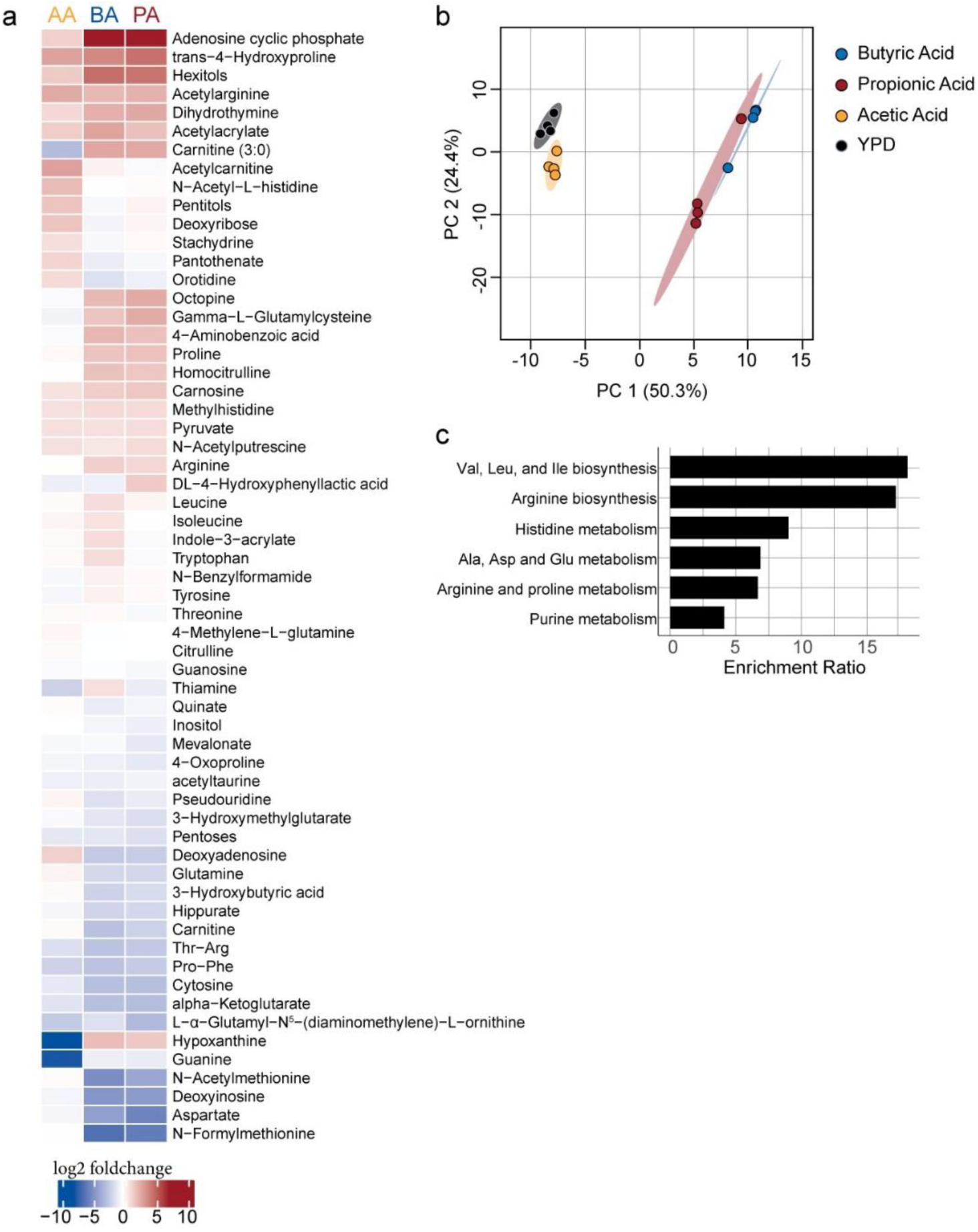
SCFAs disrupt central carbon and amino acid metabolism in *C. albicans*. (a–b) Intracellular metabolites of SC5314 cells cultured ±50 mM butyrate (BA), propionate (PA), or acetate (AA) for 30 min under aerobic conditions were profiled by hydrophilic interaction liquid chromatography–mass spectrometry (HILIC–MS). Of 591 metabolites screened, 148 were detected. **(a)** Principal component analysis (PCA) showing metabolic signatures after BA, PA, or AA treatment. **(b)** Heatmap of metabolites significantly altered by BA, PA, or AA compared with untreated controls. **(c)** KEGG pathway enrichment analysis of differentially abundant metabolites. Data represent mean ± SEM from three independent experiments. Differential expression analysis was performed using the limma package, and significance for metabolite differences was determined using false discovery rate (FDR)–adjusted *p* < 0.05.

Metabolic pathway enrichment analysis revealed broad suppression of amino acid biosynthesis and metabolism pathways and nucleotide metabolism (**Fig. 3c**; **Supplemental Tables, “Metabolomics_enrich” tab**), consistent with inhibition of central carbon metabolism. Taken together with the transcriptional and glucose uptake data, these results indicate that SCFAs impair *Ca* growth by constraining glucose entry, depleting TCA cycle intermediates, and suppressing amino acid and nucleotide biosynthetic pathways. BA and PA imposed nearly identical metabolic constraints, which were distinct from the profile induced by AA.

### SCFAs induce *Ca* intracellular acidification

SCFAs inhibit bacterial growth in part through intracellular acidification^9,10^. Notably, intracellular pH is closely linked to glucose sensing and uptake in *Ca* and *Saccharomyces cerevisiae*^55,56^. In *S. cerevisiae,* low extracellular glucose reduces intracellular pH to 6.2 - 6.4^55,56^. Given these observations and the SCFA-mediated downregulation of *Ca* glucose sensing gene *GPR1* (**Fig. 2c**), we hypothesized that SCFAs lower *Ca* intracellular pH.

To test this, we used *Ca* strain JKC1559^57^, which expresses a pH-sensitive GFP variant (the local pH changes its fluorescence spectra^58,59^), to directly measure *Ca* intracellular pH levels (**Fig. 4a**). BA and PA (10 mM) significantly reduced *Ca* intracellular pH compared to untreated controls. The combination of all three SCFAs produced an intermediate, but significant decrease, and AA required 25 mM to elicit a smaller reduction (**Fig. 4b**).

**Figure 4.**
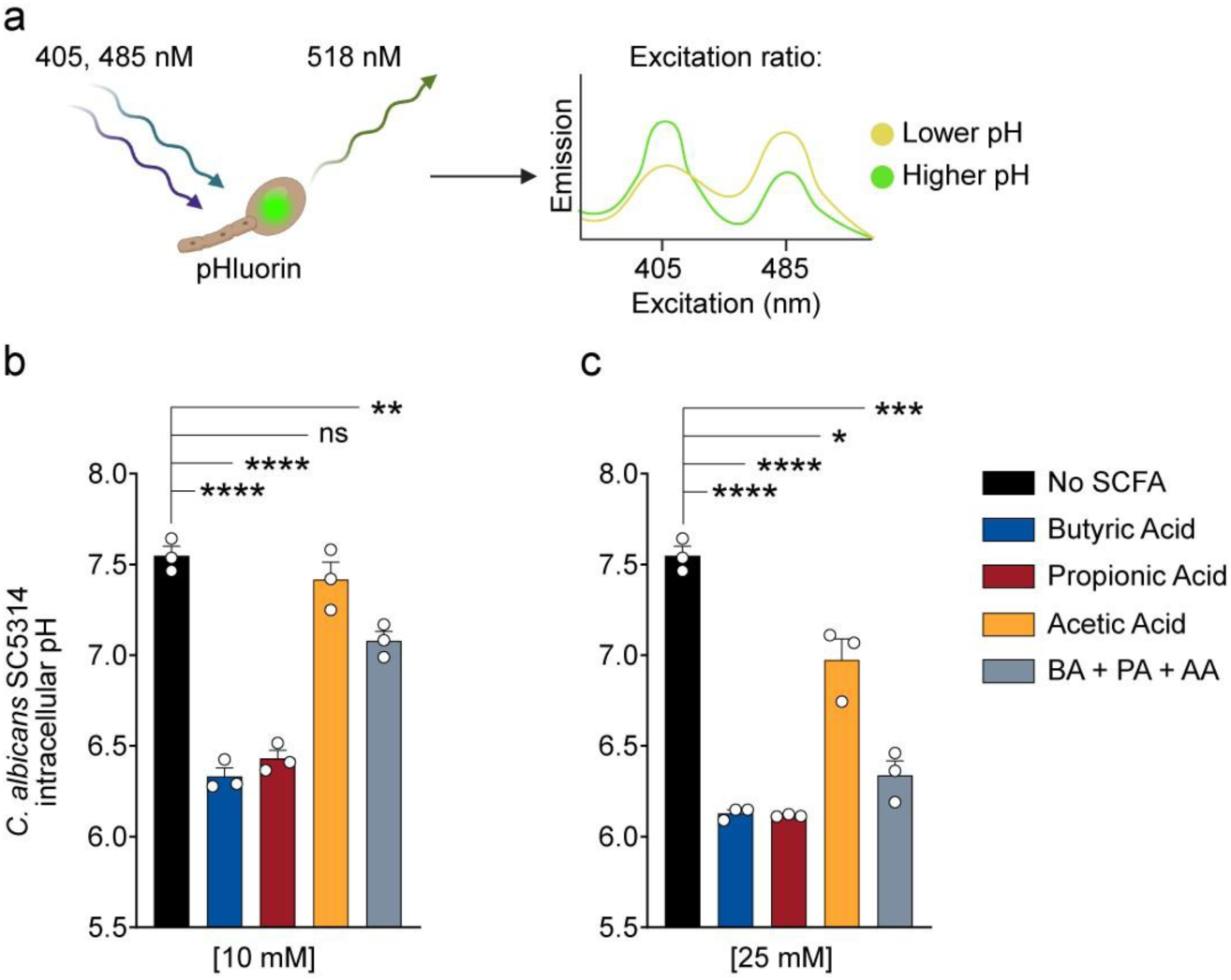
SCFAs induce intracellular acidification in *C. albicans*. **(a)** Schematic of intracellular pH measurement using strain JKC1559 expressing the pH-sensitive GFP variant pHluorin. Intracellular pH calculated from the ratio of emission at 518 nm following excitation at 405 and 485 nm, based on a calibration curve generated from cells incubated in media of defined pH. **(b–c)** Intracellular pH of JKC1559 cells grown at 30°C in YNB LO-FLO medium and incubated for 10 min + SCFA. Media pH was adjusted to 5.0 across conditions. **(b)** Intracellular pH after 10 mM SCFA treatment. **(c)** Intracellular pH after 25 mM SCFA treatment. Bars = mean ± SEM from three independent experiments. Unpaired *t*-test: \**P* < 0.05, \*\**P* < 0.01, \*\*\**P* < 0.001, \*\*\*\**P* < 0.0001, ns, not significant.

In summary, BA and PA are potent inducers of intracellular acidification in *Ca*. This effect likely contributes to growth inhibition by impairing metabolic enzymes, depleting energy, and disrupting stress-response signaling. Together with repression of central carbon metabolism (**Figs. 2, 3**), intracellular acidification appears to be a central mechanism of SCFA-mediated growth suppression.

### BA and PA reduce *Ca* colonization in the murine GI tract

To determine whether SCFAs reduce *Ca* colonization *in vivo*, we adapted a murine model of *Ca* gastrointestinal colonization^15-17,45^. Briefly, C57BL/6 mice were pre-treated with antibiotics to deplete endogenous commensal gut bacteria and reduce baseline GI SCFA levels (**Fig. 1b**) and then colonized with *Ca*. Mice were supplied sterile drinking water supplemented with SCFA (150 mM AA, BA, PA or combination SCFA; pH 5.0^7^) (**Fig. 5a**).

**Figure 5.**
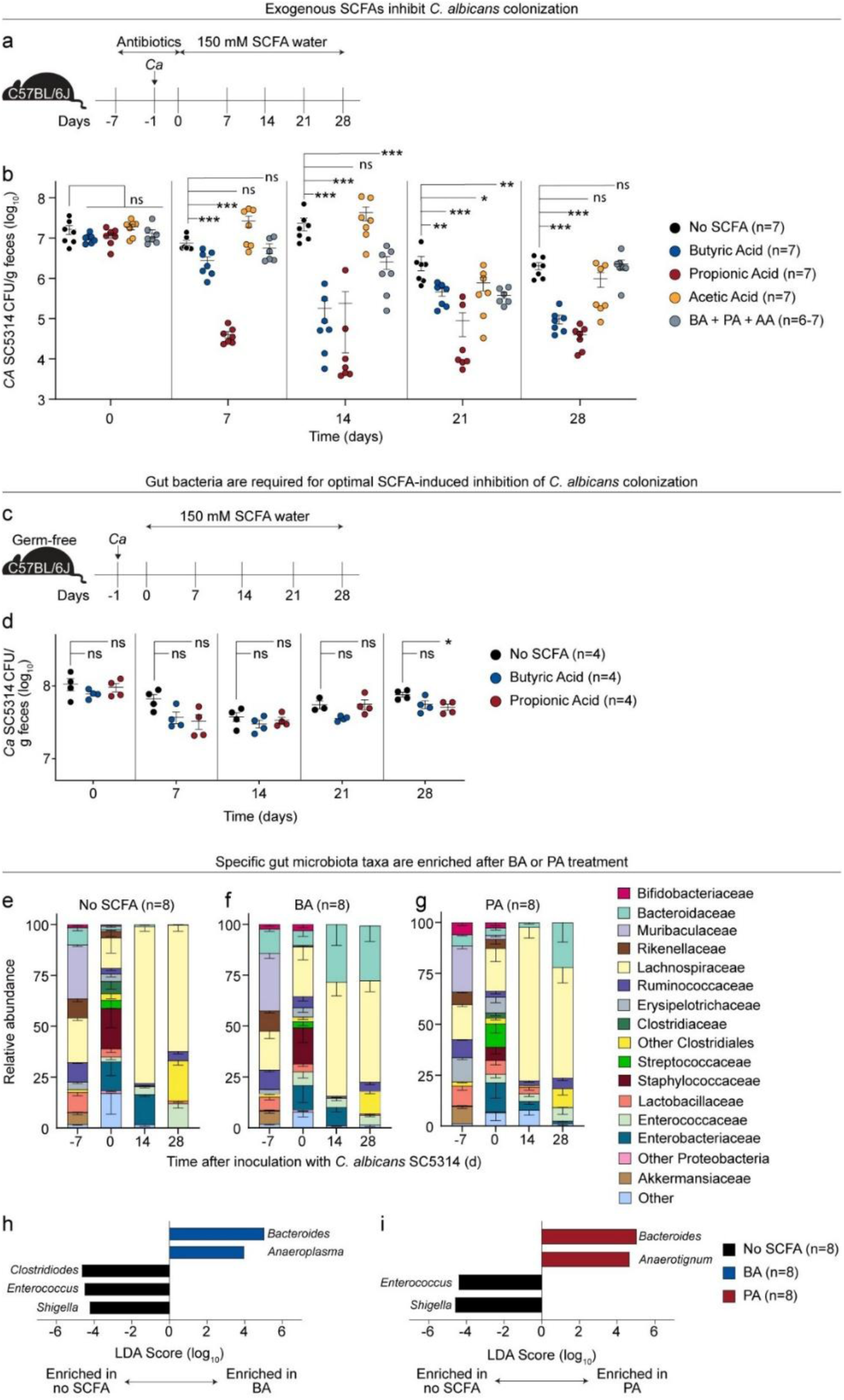
BA and PA reduce *C. albicans* colonization in the murine gastrointestinal tract. **(a)** Experimental schema for assessing SCFA treatment in conventional C57BL/6J mice (female, 6–8 weeks, Jackson). Mice received penicillin/streptomycin in drinking water for 7 days to deplete commensals, were orally gavaged with 2 × 10^8^ CFU *C. albicans* SC5314, and then switched to sterile, pH-adjusted (5.0) drinking water ±150 mM SCFA ad libitum. Colonization was monitored for 28 days. **(b)** *Ca* GI colonization levels, as determined by cultured enumeration of fecal homogenates on YPD with gentamicin and vancomycin. Data from two independent experiments; n = 6–7 mice per group. **(c)** Experimental schema for SCFA treatment in germ-free C57BL/6 mice (female, 6-10 weeks, Gnotobiotic facility, UTSW). Mice were orally gavaged with 2×10^8^ CFU *C. albicans* SC5314 and treated with sterile, pH-adjusted water ±150 mM SCFA ad libitum. GI Colonization measured for 28 days. **(d)** *Ca* GI colonization levels, determined as in (**b**). Data from two independent experiments; n = 4 mice per group. **(e–g)** Gut microbiota recovery in conventional mice from (**a**), profiled by 16S rRNA gene sequencing (V4 region) of fecal gDNA. Relative abundances of bacterial families over 28 days in (e) no SCFA, (**f**) BA, and (**g**) PA groups. **(h–i)** Differential abundance of bacterial genera on day 28 comparing **(h)** BA or **(i)** PA versus no SCFA controls, determined by LEfSe analysis. Taxa with >2 log LDA score and *p* < 0.05 (Kruskal–Wallis test) are shown. Points = results from individual animals. Bars = mean ± SEM. Mann–Whitney test. \**P* < 0.05, \*\**P* < 0.01, \*\*\**P* < 0.001, ns, not significant.

Mice treated with BA or PA exhibited significantly lower *Ca* GI colonization levels, as early as day 7, compared to controls receiving sterile acidified water (**Fig. 5b**). In contrast, AA or SCFA combination treatments did not result in significant reductions in *Ca* GI levels until days 21 or 14, respectively (**Fig. 5b**). Overall, BA and PA resulted in the lowest *Ca* GI colonization levels across the entire 28-day experiment, findings consistent with our *in vitro* findings (**Fig. 1**), in which AA was the least effective in inhibiting *Ca* growth.

Inbred mouse strains can have distinct gut bacterial microbiota and phenotypes^60-62^. To exclude mouse strain-specific effects, we repeated the experiment using Envigo C3H/HeN mice. Consistent with our results in Jackson C57BL/6 mice, only BA and PA significantly reduced *Ca* distal gut colonization levels; AA or combination SCFAs had no effect (**Fig. S3**). BA or PA treatments also consistently lowered *Ca* luminal burden in proximal GI segments (duodenum, ileum, cecum) and colon, compared to water treatment (**Fig. S4**).

Overall, these data indicate that exogenous SCFA administration – particularly with BA and PA – reduces *Ca* colonization in antibiotic-treated mice. Thus, reintroducing SCFAs may restore functional colonization resistance in the antibiotic-disrupted gut and limit fungal overgrowth.

### Gut microbiota are essential for SCFA-mediated protection against *Ca* GI colonization

To test whether SCFAs alone, in the absence of gut microbiota, could reduce *Ca* GI colonization, we administered SCFAs to *Ca-*colonized germ-free mice (**Fig. 5c**). BA or PA treatment produced only modest reductions in *Ca* GI levels (∼0.2-0.25 log) (**Fig. 5d**), in contrast to the 2-2.5 log fold reduction in conventional mice treated with SCFAs (**Fig. 5b**). SCFAs are largely absorbed in the proximal gut^63-65^ and may not reach concentrations sufficient to directly inhibit *Ca* growth/colonization in the distal intestine.

Because cessation of antibiotics in this murine model permits gut microbiota recovery sufficient to restore colonization resistance ^5,15^, we next asked whether SCFA efficacy depends on microbiota reconstitution. Additionally, SCFAs can promote the recovery and enrichment of beneficial gut taxa in mice and humans^66,67^. In conventional mice maintained on continuous antibiotics, BA and PA reduced *Ca* colonization only modestly (∼0.5 log at days 14 and 28; **Fig. S5**), in stark contrast to the marked protection observed when antibiotics were discontinued (**Fig. 5b**).

Gut microbiome profiling revealed distinct taxonomic shifts during recovery across treatment groups (**Fig. 5e-g**). Specifically, *Bacteroides* were significantly enriched in both BA- and PA-treated mice at day 28 post SCFA treatment, (linear discriminate analysis coupled with effect size measurements, LEfSe; p= 0.0063 for BA; p= 0.0117 for PA; Kruskal-Wallis test; **Fig 5h, i**). BA treatment led to enrichment of *Anaeroplasma*, a low-abundance commensal linked to reduced inflammation and improved intestinal barrier function in mice^68,69^. Finally, *Anaerotignum* (Lachnospiraceae), a known producer of acetate and propionate^70-72^, was significantly enriched in the PA treated group (LEfSe, p= 0.0038). In summary, SCFAs alone are insufficient to reduce *Ca* colonization in the absence of gut microbiota. Optimal protection requires microbial recovery, with BA and PA promoting enrichment of taxa that may contribute to re-establishing colonization resistance.

### SCFA protection persists in immune- and receptor-deficient mice

SCFAs exert broad immunomodulatory effects, including inducing antimicrobial peptide secretion. CRAMP, an antimicrobial peptide effective against *Ca*, is regulated by the transcription factor, hypoxia inducible factor 1α (HIF-1α), in myeloid and epithelial cells^73^. Previously, we demonstrated that pharmacologic activation of HIF-1α augments CRAMP production in the gut and reduces *Ca* GI colonization and dissemination in mice^5^.

To test whether SCFAs modulate HIF-1α/Hif1α and LL-37/CRAMP, we exposed human and mouse colonocytes to SCFAs. Indeed, SCFA treatment significantly increased expression of both HIF-1α and CRAMP in cultured human colonocytes (**Fig. S6a**) and mouse colon tissue (**Fig. S6b**). To assess the effect of these immune effectors *in vivo*, we treated *Camp* knockout and intestinal epithelial cell-specific *Hif1α* knockout mice with SCFAs. In both models, BA and PA significantly reduced *Ca* colonization (**Fig. S7**), suggesting that loss of CRAMP or HIF-1α alone is not sufficient to abrogate SCFA-mediated protection against *Ca* GI colonization.

We next investigated whether SCFA sensing via surface and intracellular receptors mediates protection against *Ca* colonization. The major surface receptors, FFAR2 and FFAR3, modulate antimicrobial peptide expression and T_reg_ expansion^7,8,74-80^. Butyrate also activates the intracellular nuclear receptor PPARγ^11,20^. BA promotes oxidative phosphorylation in intestinal epithelial cells through PPARγ, decreasing luminal oxygen concentration^21,81-83^ and suppressing the expansion of facultative anaerobes^11,20,21^, including *Ca*^12^. To determine whether loss of individual SCFA receptors impairs SCFA-mediated protection, we assessed BA and PA treatment in *FFAR2*^-/-^, *FFAR3^-/-^*, and intestinal epithelial-specific *PPARγ* knockout mice. In all three receptor-deficient models, BA and PA still significantly reduced *Ca* colonization (**Fig. S8**). Notably, FFAR2 and FFAR3 share SCFA ligands but differ in binding affinities and downstream signaling pathways^84-86^. These results suggest that no single SCFA receptor is essential, and functional redundancy among SCFA-sensing pathways likely preserves antifungal protection.

### Modulating gut propionate alters *C. albicans* GI colonization in mice

Colonization of germ-free mice with *B. theta* reduces *Ca* colonization by ∼ 5 log fold ^5^. *B. theta* produces both AA and PA, but not BA. Given PA’s greater inhibitory effect on inhibiting *Ca* growth and colonization, we hypothesized that *B. theta*-mediated suppression of *Ca* may depend on its ability to produce PA.

To test this, we utilized a PA-deficient *B. theta* mutant (*Δprop*, lacking genes for propionate synthesis)^9,87^ and compared it to a wildtype *B. theta Δtdk* strain. Germ-free mice were first colonized with *Ca* strain SC5314, followed by co-colonization with either the wildtype or *Δprop B. theta* strain (**Fig. 6a**). Indeed, mice co-colonized with *Ca* and wildtype *B. theta* exhibited significantly lower *Ca* levels than counterparts co-colonized with the *Δprop* mutant (**Fig. 6b**). Further, cecal PA concentrations were significantly lower in mice colonized with the *Δprop* mutant compared to the wildtype *B. theta*-colonized mice (**Fig. 6c**).

**Figure 6.**
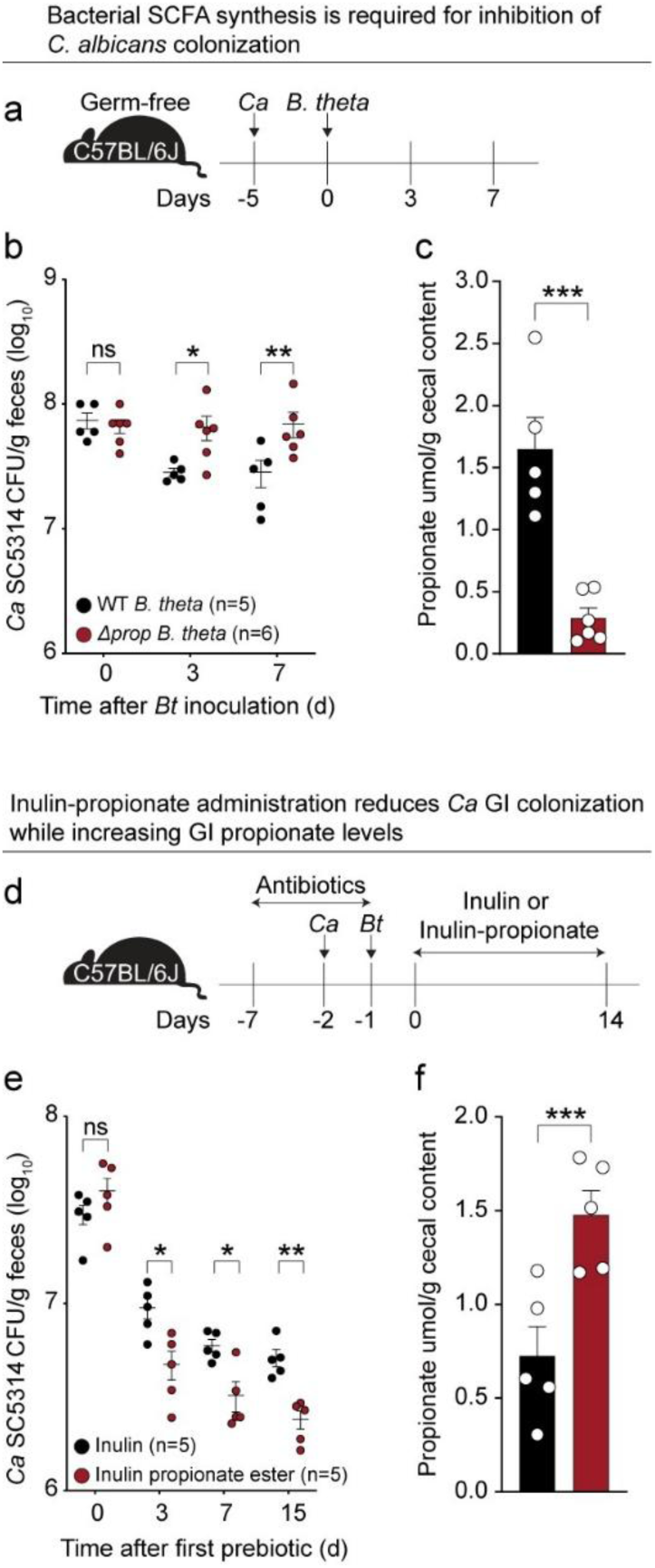
Modulating gut propionate levels alters *Ca* gastrointestinal colonization in mice. **(a)** Experimental schema for assessing the role of *Bacteroides thetaiotaomicron* (*B. theta*)–derived propionate in germ-free mice C57BL/6 mice (female, 6-10 weeks, Gnotobiotic facility, UTSW). Mice were first colonized with *Ca.* SC5314, then 5 days later with 2×10^8^ CFU either wild-type *B. theta* (Δ*tdk*) or a propionate-deficient mutant (Δ*prop*). **(b)** GI colonization levels of *Ca*, determined by cultured enumeration of fecal homogenates on YPD with gentamicin and vancomycin. Data from two independent experiments; n = 5–6 mice per group. **(c)** Cecal propionate concentrations from mice in (**a**), quantified by GC–MS. **(d)** Experimental schema for assessing the effect of inulin propionate ester in conventional C57BL/6J (female, 6-8 weeks, Jackson) mice. Mice were pre-treated with antibiotics, colonized with *Ca* SC5314, and then colonized wild-type *B. theta*. Mice were treated daily by oral gavage with 200 µl of 1% inulin or inulin propionate ester. **(e)** GI colonization levels of *Ca*, determined as in (**b**). Data from two independent experiments; n = 5 mice per group. **(f)** Cecal propionate concentrations from mice in (**d**), quantified by GC–MS. Each point = data from an individual animal. Bars = mean ± SEM. Mann–Whitney or unpaired t-test (as indicated by normality evaluation). \**P* < 0.05, \*\**P* < 0.01, \*\*\**P* < 0.001.

Prebiotics, dietary substrates selectively utilized by gut microbiota, can foster growth or activity of beneficial taxa^88^ and increase gut SCFA production^6^. Indigestible starches, such as inulin, can increase endogenous SCFA levels. To determine whether targeted prebiotic intervention could augment gut PA levels and decrease *Ca* colonization, we utilized an inulin propionate ester, synthesized by attaching a propionate moiety onto inulin. The inulin propionate ester can only be digested by specific gut microbiota residing in the distal gut (e.g. *B. theta*), directly increasing the levels of PA in the distal GI tract^9^. *Ca*-colonized mice were orally gavaged with wildtype *B. theta* (VPI-5482), then treated daily with either inulin or inulin propionate ester for 14 days (**Fig. 6d**). Mice receiving inulin propionate ester exhibited significantly lower fungal loads (**Fig. 6e**) and exhibited higher cecal PA levels when compared to inulin only treated groups (**Fig. 6f**). Overall, these results demonstrate that microbial propionate production is required for optimal suppression of *Ca* colonization in the murine gut. Prebiotic augmentation of luminal propionate further reduced fungal burden, underscoring the potential for microbiota-targeted strategies to enhance colonization resistance.

## Discussion

This study defines a direct, mechanistically supported role for microbiota-derived SCFAs in restricting *Ca* GI colonization. We show that SCFAs produced by anaerobic commensals inhibit *Ca* growth through a multifaceted mechanism involving intracellular acidification, impairment of hexose uptake, and disruption of central carbon metabolism. These effects are conserved across clinical isolates and are more potent under anaerobic conditions, emphasizing their physiological relevance in the gut environment.

While bacterial colonization resistance has been well documented, the mechanisms of fungal colonization resistance remain comparatively underexplored. Previous studies have correlated antibiotic-induced SCFA depletion with increased *Candida* colonization in mice^5,89^. Yasume-Mitobe et al. (Hohl lab, co-submitted) extend these observations to human stem cell transplant patients, further validating the inverse relationship between gut SCFA levels and fungal burden. Several gut commensals^90-93^—including *Bacteroides thetaiotaomicron* and *Blautia producta ^4,5^*—have been implicated in promoting colonization resistance and share the capacity to produce SCFAs, suggesting that metabolite output underpins their antifungal effects. Together, our studies highlight convergent metabolite-based mechanisms of colonization resistance: their work identifies valerate as a potent inhibitor of *C. parapsilosis*, while ours defines a broad SCFA program that constrains *C. albicans*. Importantly, both studies converge on intracellular acidification as a unifying antifungal mechanism, underscoring a principle of trans-kingdom ecology by which commensal bacteria limit fungal expansion.

At the mechanistic level, our transcriptomic and metabolomic analyses revealed that BA and PA suppress key metabolic programs in *Ca,* including glucose sensing (*GPR1*), hexose transport (*HGT1*), and carbohydrate metabolism (*TYE7*). These effects are consistent with prior studies linking SCFAs to repression of ribosomal biogenesis^94^ and the glucose transporter expression (*HGT16)^95^*. We extend these findings by demonstrating SCFA-induced reductions in glucose uptake and broad metabolic derangement, including depletion of tricarboxylic acid cycle intermediates and broad suppression of amino acid and nucleotide metabolism. Together with intracellular acidification, these findings suggest that SCFAs drive *C. albicans* into a starvation-like state incompatible with sustained growth.

Our results also clarify the dependence of SCFA protection on an intact gut microbiota. Germ-free or continuously antibiotic-treated mice showed only modest reductions in colonization following SCFA administration, in stark contrast to the robust protection observed in conventional mice. This indicates that exogenous SCFAs alone are insufficient to suppress fungal colonization and that cooperative microbial recovery is required. Indeed, we observed BA- and PA-driven enrichment of taxa such as *Bacteroides*, *Anaeroplasma*, and *Anaerotignum*, which may contribute to re-establishing colonization resistance. These findings highlight a dual role for SCFAs: direct inhibition of fungal metabolism and indirect promotion of protective microbial consortia.

Interestingly, SCFA-mediated protection persisted in mice lacking individual immune effectors (CRAMP, HIF-1α) or SCFA receptors (FFAR2, FFAR3, PPARγ), suggesting that direct fungal inhibition and/or redundancy among host pathways may be sufficient to preserve antifungal effects. This does not diminish the important contributions of immune and epithelial cell modulation, but it does emphasize that in our model the antifungal activity of SCFAs can be maintained in the absence of single host factors.

Several important questions remain. The intracellular pathways linking acidification to metabolic arrest are incompletely defined and may involve redox stress, protein denaturation, or epigenetic remodeling. SCFAs are known histone deacetylase inhibitors^96^, and crotonate—a less abundant SCFA—has been shown to regulate *C. albicans* hyphal gene expression and reduce invasive growth^97^. Future studies should dissect how these epigenetic effects intersect with metabolic regulation, stress signaling and morphogenetic programs. Further, the intricate interplay between bacterial producers, SCFA availability in the gut, gut environmental conditions (e.g. oxygen concentrations) and host physiology (including absorption dynamics and host immune responses) warrants further study to clarify how SCFA gradients shape fungal ecology *in viv*o.

In conclusion, our findings support a model in which SCFAs orchestrate a multifaceted antifungal program: directly suppressing fungal metabolism, shaping the gut microbiota toward a protective configuration, and potentially reinforcing epithelial and immune defenses. Together with the companion study by Yasume-Mitobe et al., these data position SCFAs as central mediators of fungal colonization resistance across species and metabolites. The translational implications are clear: strategies to augment SCFA levels (via diet, engineered probiotics, or SCFA-conjugated prebiotics) represent a promising avenue to restore colonization resistance in patients at high risk for invasive candidiasis.

## Acknowledgments

We thank Julia Koehler, MD for providing us with the pHluorin expressing *Candida albicans* strain used in the study. We thank Denise Monack, PhD for providing us with *Δprop B.theta* mutant and *Δtdk B.theta* strain used in the study. We thank The Children’s Research Institute Metabolomics Core, UTSW for performing HILIC metabolomics profiling.

## Author contributions

A.A.M, L.V.H and A.Y.K. designed the research. A.A.M., C.M.Z., L.A.C., N.P. performed experiments. A.A.M, C.M.Z., S.G., J.K., X.Z., and A.Y.K. analyzed data. L.V.H provided with gnotobiotic mouse and support. S.E.W helped with short-chain fatty acid measurements and shared PPARγ conditional KO breeders. A.A.M, C.M.Z. and A.Y.K. edited the manuscript. A.A.M, C.M.Z. and A.Y.K. wrote the paper. All authors revised the manuscript and approved its final version.

## Data and materials availability

The 16S rRNA and RNASeq data for this study will be deposited in the National Center for Biotechnology Information (NCBI) Sequence Read Archive: http://ncbi.nlm.nih.gov/sra and http://ncbi.nlm.nih.gov/sra/. Data deposition is ongoing. Requests for microbial strains should be addressed to A.Y.K. and should be covered by a material transfer agreement. All genetically engineered mice used in this study are commercially available. All data needed to evaluate the conclusions in the paper are present in the paper or the Supplementary Materials.

## Material and Methods

### Growth of *C. albicans* isolates

Unless otherwise specified, isolates (SC5314, Can092, or CHN1) were cultured aerobically overnight in 30 ml YPD (10g/L yeast extract, 20g/L peptone, 2% dextrose) at 30°C. Cultures were then harvested, washed twice with sterile PBS and resuspended in 10 ml PBS for further processing, according to the experiment. YVG plates (20g/L agar, 30mg/L vancomycin, 30mg/L gentamicin in YPD) were used to grow and quantify *Ca* isolates from *in vitro* cultures and *in vivo* fecal or GI luminal homogenates. Vancomycin and gentamicin are used to prevent gram-positive and gram-negative bacterial growth, respectively. Cultures or homogenates are serially diluted and 10 µl of dilution is plated for enumeration.

### Aerobic growth curve of *C. albicans* with short-chain fatty acids (SCFA)

*Ca* was grown aerobically overnight in YPD, harvested, washed twice with sterile PBS, and resuspended in 10 ml PBS. For aerobic growth curve experiments, isolates were seeded in YPD in 96-well plates at a starting OD_600_ of 0.1. The wells had varying concentrations of SCFA (butyric acid, Sigma-Aldrich, B10350; propionic acid, Sigma Aldrich, P1086; or acetic acid, Amresco, 0714) or HCl (Fischer, 414) in YPD. Cells were grown at 37°C with orbital shaking in a plate reader (Biotek Synergy HT) for 12 hours. OD_600_ reading was taken every hour to monitor growth in each well. Experiments were performed independently three times, with each experiment having three technical replicates. Growth curves were compared using the linear mixed effect model with Kruskal-Wallis test for significance.

### Anaerobic growth curve of *C. albicans* with SCFA

*Ca* was grown aerobically overnight in YPD and processed as above. Isolates were seeded in YPD in 96-well plates at a starting OD_600_ of 0.1. Individual wells had varying concentrations of SCFA or HCl in YPD. Cultures were grown at 37°C without shaking for 96 hours in an anaerobic chamber (Coy). OD_600_ reading was taken every 24 hours to monitor growth in each well. Experiments were performed independently three times, with each experiment having three technical replicates. The curves were compared using the linear mixed effect model with Kruskal-Wallis test for significance.

### Measuring anaerobic growth of *C. albicans* using dilution plating with SCFA

*Ca* was grown aerobically overnight in YPD and processed as above. Isolates were seeded in YPD in 12-well plates at a starting concentration of 10^5^ cells/ml. Each well contained 25 mM of SCFA at pH 5.0 or YPD, pH 5.0 (adjusted with HCl). Cells were grown at 37°C without shaking for 4 hours in an anaerobic chamber at 37°C. Cultures from each well were serially diluted and plated on YPD plates with vancomycin and gentamicin (YVG) plates which were then incubated overnight at 30°C, aerobically. Colonies were counted to evaluate the growth. Experiments were independently performed two times, with each experiment having three technical replicates. Differences in growth were compared using unpaired parametric t-tests.

### in vitro Candida albicans RNASeq Experiment

*C. albicans* strain SC5314 was grown in YPD with or without 50 mM acetic acid, 50 mM butyric acid, or 50 mM propionic acid. pH was adjusted to 5.0 across all samples. Cultures were grown in aerobic conditions at 37°C for 30 minutes. *Ca* cells were harvested, washed with sterile PBS thrice, and flash frozen. Total RNA was extracted using Trizol-chloroform extraction. Crude RNA was treated with DNaseI (Qiagen) and column purified (RNEasy Kit, Qiagen). RNA concentrations were quantified by spectrophotometry (Nanodrop, ThermoScientific). The quality of the resultant RNA was determined using an Agilent Bioanalyzer (Agilent). For samples with RNA Integrity Numbers of greater than 7.5, the RiboMinus Kit (Life Technologies) was used to deplete rRNA from the total RNA samples. 500 ng of mRNA was used to create cDNA libraries (UTSW Microarray Core). Paired-end libraries for Illumina sequencing were prepared from purified cDNA (TruSeq RNA Sample preparation kit, Illumina). Sequencing was performed on an Illumina Hi-Seq (PE-150). Reads were aligned to *Ca* SC5314 genome using CLC-Biosystems RNA-Seq module. For each ORF, the number of reads per kilobase of gene model per million mapped reads (RPKM) was calculated ^98^. The raw read counts for each gene were processed using DESeq software (utilizing negative binomial distribution analysis). Significantly differentially expressed genes were defined as ≥ 2 fold-change and adjusted p-values < 0.05 (Benjamini-Hochberg procedure)^99^, when compared to the untreated control group. KEGG (Kyoto Encyclopedia of Genes and Genomes) analysis was performed for the shared downregulated genes across BA, PA and AA treatment to identify effected pathways. Gene ontology analysis was performed using the Go Term Finder tool in Candida Genome database (http://www.candidagenome.org/cgi-bin/GO/goTermFinder) in August 2025. Significance was determined by Bonferroni correction of p-values, and only terms with an adjusted p-value ≤ 0.05 were included.

### 2-deoxyglucose uptake assay

Glucose uptake assay kit from Abcam was used (Product # 136956). SC5314 was grown aerobically overnight in YPD at 30^ο^C. Cells were harvested the next day, washed with PBS three times and resuspended at a concentration of 10^8^ cells/ml. Cells were then pelleted and resuspended in Krebs-Ringer-Phosphate-Hepes buffer (20 mM Hepes, 5 mM KH2PO4, 1 mM MgSO4, 1 mM CaCl2, 136 mM NaCl, 4.7 mM KCl, pH 7.4) at the same concentration, followed by incubation at 37^ο^C for 120 minutes to starve the cells of glucose. 100 µl of the starved cells were then aliquoted for each technical replicate in each group. The cells were spun down and resuspended in 100 µl YP (10 g/l yeast extract and 20 g/l peptone). The cells then received varying concentrations of SCFA and were incubated for 20 minutes. After 20 minutes, 10 µl of 10 mM 2-DG was added to each replicate and incubated for another 20 minutes. 10 µl of each sample was plated to quantify the viability in each sample. The cells were then washed three times with sterile PBS, and 90 µl of extraction buffer was added. Samples were flash frozen, thawed, and incubated at 85°C for 40 minutes. The cells were then cooled and 10 µl of neutralization buffer was added. The contents were mixed thoroughly and centrifuged for 2 minutes at 500g, room temperature. 50 µl of supernatant were transferred to black 96-well plates with clear bottoms. To generate the standard curve, 50 µl of six 2-DG standards were also added in duplicate. 50 µl of reaction mix was added to all the samples and standards and incubated at 37°C for 40 minutes. The plate was then measured (excitation wavelength of 537 nm, emission at 585 nm) in a plate reader (Synergy HT). The readings were then converted into 2-DG concentration based on the standard curve. The concentration was then normalized by the cell viability of each sample to quantify the 2-DG uptake. Unpaired parametric t-tests were used to analyze the significance of the difference in 2-DG uptake between different groups.

### Intracellular pH measurement in *C. albicans*

To evaluate the intracellular pH in *C. albicans*, strain JKC1559 (which has the pHluorin gene encoded in its genome and was generously provided by Julia Kohler, MD, Harvard Medical School) was utilized. A calibration curve was generated by the following protocol: JKC1559 was grown overnight in YPD for 19 h. The cells were washed twice with 30 ml normal saline (NS). Cells were centrifuged at 2000g for the washes. The washed cells were weighed and resuspended in NS at a concentration of 0.5 g/ml. 100 µl of cells were then inoculated into 10 ml of the YNB LO FLO (Formedium, CYN6202) medium and incubated at 30°C, aerobically for 4 hours. Cells were then washed with the culturing medium once and resuspended to a final concentration of 0.5 g/ml cells. 20 µl of cell suspension was added to each pH calibration tube containing 2 ml of YNB LO FLO medium at specific pHs (pH 5.0, pH 5.5, pH 6.0, pH 6.5, pH 7.0, pH 7.5). Additionally, 20 µl of *Ca* suspension was added to a tube with 110 μM monensin, and 15 μM nigericin. The tubes were then incubated at 30°C for one hour, aerobically, with gentle mixing. 300 μl of each sample was transferred to 96-well plate and fluorescence was measured using a plate reader (Synergy HT) at excitation wavelength of 405 nM and 485 nM, with emission wavelength at 518 nM. Ratio of emission intensity at 405 nM and 485 nM was calculated for all the samples and plotted against the pH of each of calibration standard to generate the calibration curve. This curve was used to extrapolate the pH of the experimental samples.

For experimental samples, cells were grown and harvested (exactly as noted previously to generate the calibration curve) to the final density of 0.5 g/ml. This final culture was then used to inoculate 297 µl YNB LO-FLO media per well with varying concentrations of short-chain fatty acid in a 96-well plate. pH of media across was adjusted to 5.0. 3 µl of the culture was used to inoculate 297 µl of the media in 96-well plate. The media was mixed thoroughly and the fluorescence (405 nM and 485 nM excitation; 518 nM emission) was read 10 minutes after resuspending cells in the experimental condition. The ratio of emission intensity at 405 nM and 485 nM excitation wavelength was calculated. The calibration curve was then used to convert the ratio into corresponding intracellular pH value. Unpaired parametric t-tests were used to analyze the significance of the difference in intracellular pH between different groups.

### Short-chain fatty acid quantification

Mouse cecal contents were transferred into pre-weighed tubes, weighed, and resuspended in 1ml PBS. Samples were vortexed and centrifuged at 6000g for 15 minutes at 4°C. The supernatant was collected. 95µl of the supernatant was mixed with 5 µl of 100 µM d-butyrate. The samples were completely dried (Eppendorf Vacufuge). Dried samples were then resuspended in 100 µL of pyridine. All samples were then sonicated (water bath sonicator, 50% amplitude, Emerson) for 1 minute. The samples were then incubated for 20 minutes at 80°C. The samples were cooled and derivatized with MTBSTFA (M-108, Cerilliant). 100µL of MTBSTFA was added to all the samples and mixed thoroughly. The samples were then incubated at 80°C for one hour. After one hour, samples were centrifuged at 13,000g for 1 minute. Following centrifugation, the derivatized samples were moved to autosampler vials for gas chromatography-mass spectrometry (GC-MS) analysis (Shimadzu, TQ8040). The injection temperature was 250°C and the injection split ratio was set to 1:100 with an injection volume of 1 μl. The oven temperature was set at 50°C for 2 min, increasing to 100°C at 20°C per min and to 330°C at 40°C per min, with a final hold at 330°C for 3 min. The flow rate of the helium carrier gas (99.9999% purity) was kept constant at a linear velocity of 50 cm/s. We used a 30 m × 0.25 mm × 0.25 μm Rtx-5Sil MS (Shimadzu) column. The interface temperature was 300°C. The electron impact ion source temperature was 200°C, with 70 V ionization voltage and 150 μA current. Acetate (m/z of <u>117</u> and 159), propionate (m/z of <u>131</u>, 132, and 75), butyrate (m/z of <u>145</u>, 146, and 75) and deuterated butyrate (m/z of <u>152</u>, 153, and 76) were quantitated in single ion monitoring mode. Concentrations were calculated based on an external standard curve, with deuterated butyrate as the internal standard. Unpaired parametric t-test was used to analyze the significance in difference between propionate levels measured between groups.

### Synthesizing inulin propionate ester

Inulin propionate ester was synthesized following a previously published procedure for inulin esterification ^9^. Briefly, inulin (20 g, 0.123 mole), propionic anhydride (18.3 g, .014 mol), and 1-methyl imidazole (10.1 g, 0.123 mol) were combined in 60 ml DMSO and stirred at room temperature for 4 hours. The mixture was diluted with 120 ml water. The reaction mixture was dialyzed against water for 2 days at 4°C. Following dialysis, samples were freeze-dried. Product was tested using ^1H^NMR, a peak in 1-2 ppm region validated the successful synthesis of inulin propionate ester.

### HT-29 SCFA treatment

HT-29 cells were obtained from the ATCC (HTB-38) and cultured in DMEM (Gibco) with 10% heat-inactivated FBS (Corning, 35-016-CV) at 37°C, 5% CO_2_. For the experiment, 1.5 × 10^5^ cells were plated in each well of a 12-well plate (Falcon, Corning) and cultured for 4 days to reach confluency. Once confluent, the cells were washed twice with warm PBS and then the treatment media was added. 5 mM or 10 mM acetic acid, butyric acid, or propionic acid in DMEM, with 10% FBS were used as treatment media. For the control, DMEM, with 10% FBS was adjusted to pH 6.5 with HCl, to match the pH of the SCFA media. Each treatment was performed in triplicate. HT-29 cells were cultured with the treatment media for 4 hours at 37°C, 5% CO_2_. After the treatment, the cells were washed twice with PBS and resuspended in 1 ml Trizol (per well) (Life Technologies). The Trizol samples were collected in 500 ul aliquots and frozen at -80°C until RNA extraction. The experiment was performed three times, with different passage numbers of the HT-29 cells.

To prepare the SCFA treatment media, 10 mM stocks of each SCFA media was prepared and sterile filtered. These stocks were aliquoted and frozen at -20°C until use. On the day of the experiments, the frozen stocks were thawed at room temperature, briefly warmed to 37°C in a bead bath, then used for the experiment. The 5mM SCFA media was diluted from the 10 mM stocks with DMEM, 10% FBS, pH 6.5 media. DMEM, 10% FBS, pH 6.5 media was stored at 4°C until use or prepared fresh.

### Human colonocyte culture and mouse colon qPCR

Mouse colon was harvested and placed in tubes with 1 ml RNA*later* (Invitrogen) on ice. Colon segments in RNA*later* were incubated at 4°C overnight and stored at -80°C till RNA extraction. Colon segments stored in RNA*later* were thawed and moved to tubes containing 600 µl of RLT buffer and two 6.35 mm glass beads. Colon segments were lysed and homogenized by bead beating the samples for 1 minute, twice. Samples were centrifuged at 12000g for 3 minutes and the supernatant was moved to a new RNase-free tube. Total RNA was then isolated using the Qiagen RNEasy RNA isolation kit (Qiagen) according to the prescribed protocol.

For colonocyte HT-29 RNA extraction, samples (500 ul Trizol) were thawed on ice and 100 µl of chloroform was added to each sample, mixed and incubated at room temperature for 3 minutes. Samples were centrifuged at 12000g for 15 minutes at 4°C and 200 µl of the aqueous layer was collected in different RNase-free tube. 200 µl of 70% ethanol was added to the collected supernatant to precipitate the RNA and was further processed with the Qiagen RNeasy kit, with on-column DNase digestion, using the prescribed protocol.

Total RNA from mouse colon or HT29 cells was used to synthesize cDNA (iScript, BioRad). qPCR analysis was performed using the SsoAdvanced SYBR Green Supermix (Bio-Rad) or PowerTrack SYBR Green Master Mix (Applied Biosystems) and specific primers. Signals were normalized to 18S rRNA levels within each sample and normalized data were used to calculate relative levels of gene expression using ΔΔC_t_ analysis. Unpaired parametric t-test was used to analyze the significance in difference between relative expression level of genes between groups List of primers used are listed below.

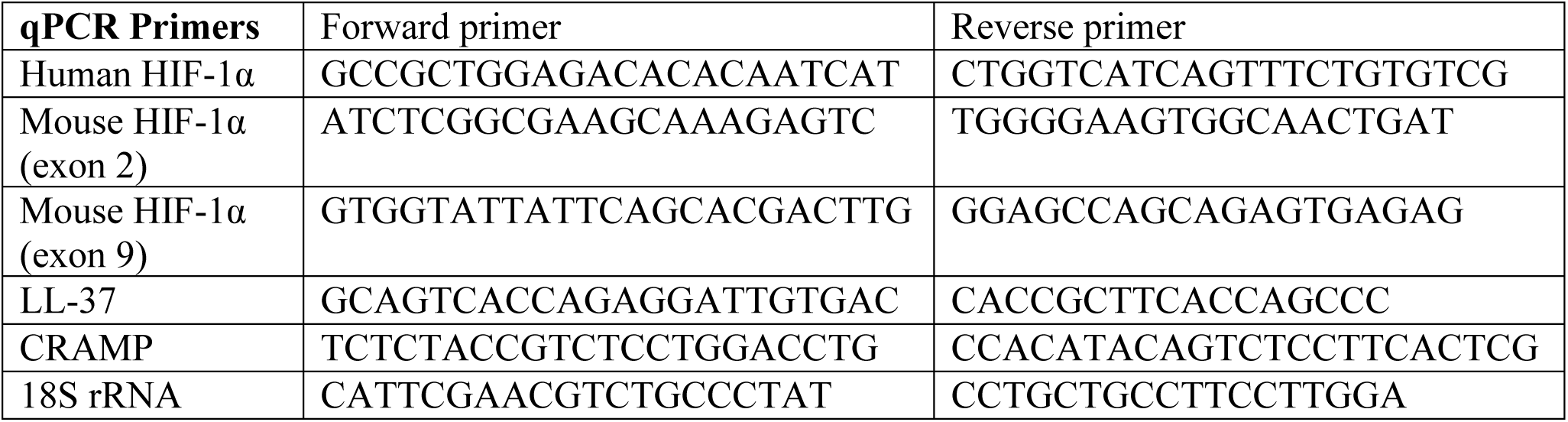

### Mouse strains used in this study

Wildtype C57BL/6J mice (Jackson, female, 6-8 weeks, stock 000664, Room RB12) and C3H/HeN mice (Envigo, female, 6-8 weeks, stock 040) were used for studying the effect of SCFA on *Ca* GI colonization. Heterozygous breeding pairs for *Ffar2* KO mice (MMRRC, stock 047690-UCD) were bred to generate *Ffar2* KO mice. Heterozygous breeding pair for *Ffar3* KO (RIKEN, stock RBRC06327) were bred to generate *Ffar3* KO mice. *Camp* KO mice (Jackson, stock 017799) were bred in our animal facility. *Hif1α^fl/fl^ vil-Cre*^+^ were bred in-house, which was originally created from B6.129-Hif1a^tm3Rsjo^/J (Jackson, stock 007561). *Pparγ^fl/fl^ vil-Cre*^+^ were propagated in-house from the breeding pair received from the laboratory of Dr. Sebastian Winter, which was originally created from B6.129-Pparg^tm2Rev^/J (Jackson, stock 004584). Germ free C57BL/6 mice were provided by the Gnotobiotic Animal Core at the University of Texas Southwestern Medical Center. Mice were fed Teklad Global 16% Protein Rodent Diet chow (Teklad 2916, irradiated). Mouse cages were changed once weekly. All the procedures performed in the experiment were done in accordance with the IACUC approved protocols.

### Effect of oral administration of SCFA on *C. albicans* SC5314 GI colonization in mice

Adult mice (female, 6-8 weeks old) were given 2 mg/ml of streptomycin and 1500 U/ml of penicillin in drinking water to deplete endogenous bacterial gut microbiota. After 7 days, bacterial microbiota clearance was checked by plating fecal pellets from mice on BHI-blood plates (anaerobic incubation, 37°C) and MacConkey and TSA plates (aerobic incubation, 37°C). Clearance was confirmed by absence of growth from these media.

*Ca* (SC5314 strain) culture was grown overnight aerobically at 30°C. Culture was harvested, washed with PBS twice and resuspended in PBS at a concentration of 1×10^9^ CFU/ml. Mice were gavaged with 200 µl of 1×10^9^ CFU/ml of *Ca*. Antibiotic treatment was stopped (after one week of administration). Mice were started either on sterile water at pH 5.0 or water with 150 mM SCFA, pH 5.0, ad libitum. Mice were fed Teklad Global 16% Protein Rodent Diet chow (Teklad 2916, irradiated). Mouse cages were changed once weekly. To quantify *Ca* distal GI tract colonization levels, mouse fecal pellets were collected, weighed, homogenized in PBS, serially diluted, plated on YVG plates and incubated aerobically at 30°C. Colonies were counted and normalized to weight of the fecal pellets to quantify *Ca* colonization levels. *Ca* colonization levels were assessed weekly for four weeks.

To determine *Ca* colonization in different GI segments, mice were euthanized on day 7 and day 28 after SCFA water initiation. Luminal contents from duodenum, ileum, cecum and colon were collected, weighed, resuspended in PBS and plated as described above for the fecal pellets.

To determine the impact of microbiota recovery on the protective effect of SCFA, the experimental scheme described above was amended so that antibiotic treatment continued for all groups, even after starting mice on SCFA water, to prevent microbiota recovery during the experiment.

In germ-free mouse experiments, no antibiotic treatment was given prior to colonizing the gnotobiotic mice with *Ca*. 6-10 weeks old female mice were used for germ-free experiments. Mice were colonized with *Ca* as described above. Mice were then put on sterile water at pH 5.0 or water with 150 mM SCFA at pH 5.0, ad libitum. Fecal pellets were sampled weekly for *Ca* levels (as described above) for four weeks after starting the groups on SCFA water. Differences in colonization levels were checked for statistical significance using Mann-Whitney tests for all experiments.

### The effect of propionate production-deficient *B. theta* mutant on *C. albicans* GI colonization

*Ca* (SC5314 strain) culture was grown overnight aerobically at 30°C. Culture was harvested, washed with PBS twice and resuspended in PBS at a concentration of 1×10^9^ CFU/ml. Germ-free mice were gavaged with 200 µl of 1 × 10^9^ CFU/ml of *Ca* strain SC5314. The following day fecal pellets were collected, weighed, homogenized in 1 ml PBS and plated on YVG plates to quantify *Ca* colonization levels. *B.theta* strains were streaked on BHI-blood plate with gentamicin (100µg/ml) in anaerobic chamber at 37°C. To grow *B.theta* cultures under anaerobic conditions, a single colony was used to inoculate TYG media with gentamicin (100µg/ml) and grown at 37°C for 48 hours in an anaerobic chamber. Cultures were then washed with reduced PBS twice and resuspended at 1×10^9^ CFU/ml. Mice were gavaged with 200µl of 1×10^9^ CFU/ml of wildtype *B.theta* (VPI5482 *Δtdk* strain) or the *B. theta Δprop* mutant strain (strain lacking the gene cluster 1686-89 and its ability to synthesize propionate). Both strains were graciously provided by Denise Monack, PhD, University of California at San Francisco. For four weeks, fecal pellets were collected every third and seventh day, and processed to quantify colonization levels of *Ca* in mice. Differences in colonization levels were checked for statistical significance using Mann-Whitney tests. Mice were euthanized at the end of the experiment, and cecal contents were collected to measure propionate levels.

### Quantifying the impact of inulin propionate ester on *C. albicans* GI colonization

Microbiota of 6-8 weeks old female C57BL/6J mice were depleted with penicillin/streptomycin treatment and colonized with *Ca* as described above. After colonization with *Ca* was confirmed, mice were put on sterile water without antibiotics. Wildtype *B.theta* cultures (VPI-5482) were grown anaerobically at 37°C as described above and resuspended in reduced PBS at a concentration of 1×10^9^ CFU/ml. Mice were gavaged with 200µl of 1×10^9^ CFU/ml wildtype *B.theta* (VPI-5482 strain). Mice were subsequently orally gavaged with 200 μL of 1% inulin or 1% inulin propionate ester daily. Fecal pellets were collected every two days, over a period of four weeks. Fecal pellets were processed to quantify the colonization levels of *Ca* in mice (as described above). Differences in colonization levels were assessed using Mann-Whitney tests. Mice were euthanized at the end of the experiment, and cecal contents were collected to measure propionate levels.

### 16S rRNA gene PCR amplification, sequencing and analysis

Bacterial gDNA was extracted from fecal samples using the MagAttract Power Microbiome DNA/RNA KF kit (Qiagen) on the Kingfisher Flex machine (Thermo Fisher Scientific). 16S rRNA genes (variable region 4, V4) were amplified from each sample in 96-well plates from respective samples^100^. We used the reverse primer 926R, 5’-CAAGCAGAAGACGGCATACGAGAT-NNNNNNNNAGTCAGTCAG-CC-GGACTACHVGGGTWTCTAAT-3”: the italicized sequence is the reverse MiSeq primer i7; NNNNNNNN designates the unique 8-base barcode used to tag each PCR product; the bold sequence is the broad-range 16S bacterial primer containing the pad-link 16SR. The forward primer used was 515F, 5’-AATGATACGGCGACCACCGAGA TCTACAC NNNNNNNN-TATGGTAATT-GT-GTGCCAGCMGCCGCGGTAA-3’: the italicized sequence is MiSeq Primer i5; the NNNNNNNN designates the unique 8-base barcode used to tag each PCR product; and the bold sequence is the broad range 16S bacterial primer containing the pad-link-16SF. PCR reactions consisted of 17ul Accuprime Pfx Supermix, 1000 nM of each primer, and 20ng of template. Reaction conditions were 2 min at 95℃, followed by 30 cycles of 20 s at 95℃, 15 s at 55℃, 5 min at 72℃, then 10 min at 72℃, and a hold at 4℃ on an Eppendorf Mastercycler. For tissue and tumor microbiome profiling, two rounds of PCR amplification were performed. Products were verified on a 1% agarose gel, and normalized using the AmPure Normalization plate protocol using the KingFisher Flex platform. Each plate was then pooled into a single tube, and the PCR product size and library quality of each individual pooled plate was checked using Agilent Technologies D1000 ScreenTape electrophoresis. Additionally, KAPA Biosystems PCR Library Quantification kit was used to quantify each pooled plate. Illumina spike in (PhiX) was included at 4 pM at 10%, and the pooled sample library was included at 4pM at 90% yielding a final library concentration of 3.6 pM and PhiX concentration of 0.4 pM. Pooled samples were then sequenced with Illumina MiSeq (PE-250) at the Microbiome Research Laboratory at UTSW Medical center.

Raw sequences generated from Illumina MiSeq for 16S rRNA gene PCR amplicons were quality filtered. Sequences shorter than 200 nucleotides or longer than 1000 nucleotides were removed. Sequences containing ambiguous bases, primer mismatches, homopolymer runs in excess of 6 bases and uncorrectable barcodes were also removed. Sequences that passed the quality filtration were de-noised and analyzed using the open-source software package Quantitative Insights Into Microbial Ecology (QIIME2). 16S rRNA gene sequences were then classified taxonomically using the classifier Silva database. Differential taxonomic abundance between different phenotypic groups was analyzed by linear discriminate analysis (LDA) coupled with effect size measurements (LEfSe). Only taxa noted to have >2 log fold increase in LDA score and p< 0.05, Kruskal-Wallis test were identified as significantly enriched or depleted.

### Metabolome extraction from *C. albicans* cells for hydrophilic interaction liquid chromatography (HILIC) analysis

*C. albicans* strain SC5314 was grown in YPD with or without 50 mM acetic acid, 50 mM butyric acid, or 50 mM propionic acid. pH was adjusted for all the samples to 5. Cultures were grown in aerobic conditions at 37°C for 4 hours. At the end of 4 hours cells were pelleted and washed with ice-cold normal saline. The metabalome was extracted as previously described^101,102^. Briefly, the pellet was resuspended in 500µl of 80% methanol and then subjected to three freeze thaw cycles in liquid nitrogen. Samples were vortexed for 1 minute and centrifuged at 20,160g for 15 minutes. Supernatant was transferred to a new tube and protein quantification was performed for all the samples using BCA protein quantification kit (Pierce;23227). Volume equivalent of 10µg of protein was transferred to another tube for each sample and completely dried in a SpeedVac. Samples were resuspended in 100 µl of 80% acetonitrile, vortexed for 1 minute and centrifuged at 20,160g for 15 minutes at 4°C. Supernatants were transferred in LC-MS vials, and HILIC metabolomics was performed^101,102^ to get the relative levels for 591 metabolites across samples. Peak assignment and metabolite level calculations were performed as described perviously^102^.

### Metabolomics Analysis

Differential expression analysis was performed using the limma package (version 3.58.1) in R (version 4.3.3). Comparisons were performed between each treatment group and the YPD control. Raw p-values were adjusted for multiple testing using the Benjamini–Hochberg procedure. Metabolites with adjusted p-values < 0.05 were considered statistically significant.

Enrichment, pathway, and principal component analyses were conducted using MetaboAnalyst 6.0 (https://www.metaboanalyst.ca/). The enrichment analysis used the KEGG metabolite set library to identify pathways significantly over-represented (Benjamini-Hochberg False Discovery Rate adjusted p-value < 0.05) among the differentially expressed metabolites. The enrichment ratio was calculated as the number of observed hits divided by the number of expected hits. Pathway analysis was performed using the hypergeometric test against the *Candida albicans* KEGG pathway library.

### Statistical analysis

GraphPad Prism v.9.2 or newer was used for statistical analysis. All datasets were tested for normality (e.g., Shapiro-Wilk). Datasets with normal distribution were analyzed with parametric tests, such as standard Student’s t-test or one-way analysis of variance (ANOVA) with Bonferroni post-test. For nonnormal distributions, nonparametric tests, such as Mann-Whitney U test or Kruskal-Wallis with Dunn’s post-test, were applied. Growth curves were analyzed using linear mixed effect model with Kruskal-Wallis test

### Language Editing Assistance

We used OpenAI’s ChatGPT (version 4.0 and 5.0) solely for language editing (e.g., grammar, clarity, and brevity) and for suggesting alternative phrasings during manuscript preparation. The tool did not contribute to study design, data generation, and analysis. It was not used to generate references. All suggestions from the ChatGPT were reviewed, revised, and approved by the authors. No text was inserted verbatim without subsequent human editing. The authors take full responsibility for the content of the manuscript.

## Supplemental Figures

**Supplemental Figure 1.**
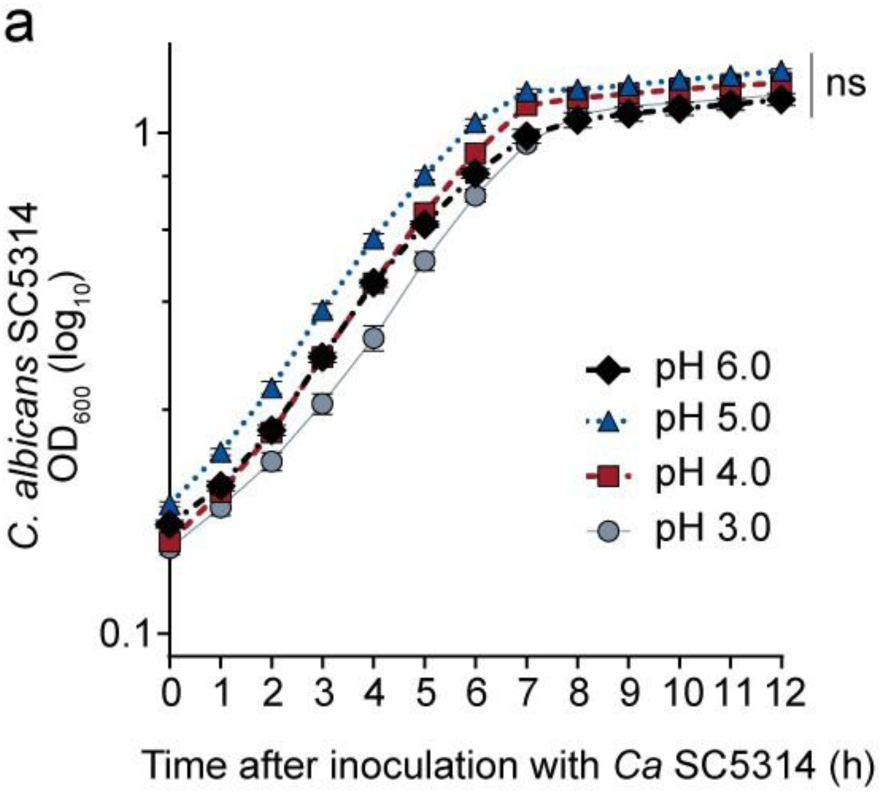
*C. albicans* growth is unaffected by low pH. **(a)** *Ca* SC5314 cultured for 12 hours in YPD at 37°C aerobically, with varying pH. OD_600_ measured every hour. Bars = mean ± SEM from three independent experiments. Linear mixed effect model with Kruskal-Wallis test. ns, not significant

**Supplemental Figure 2.**
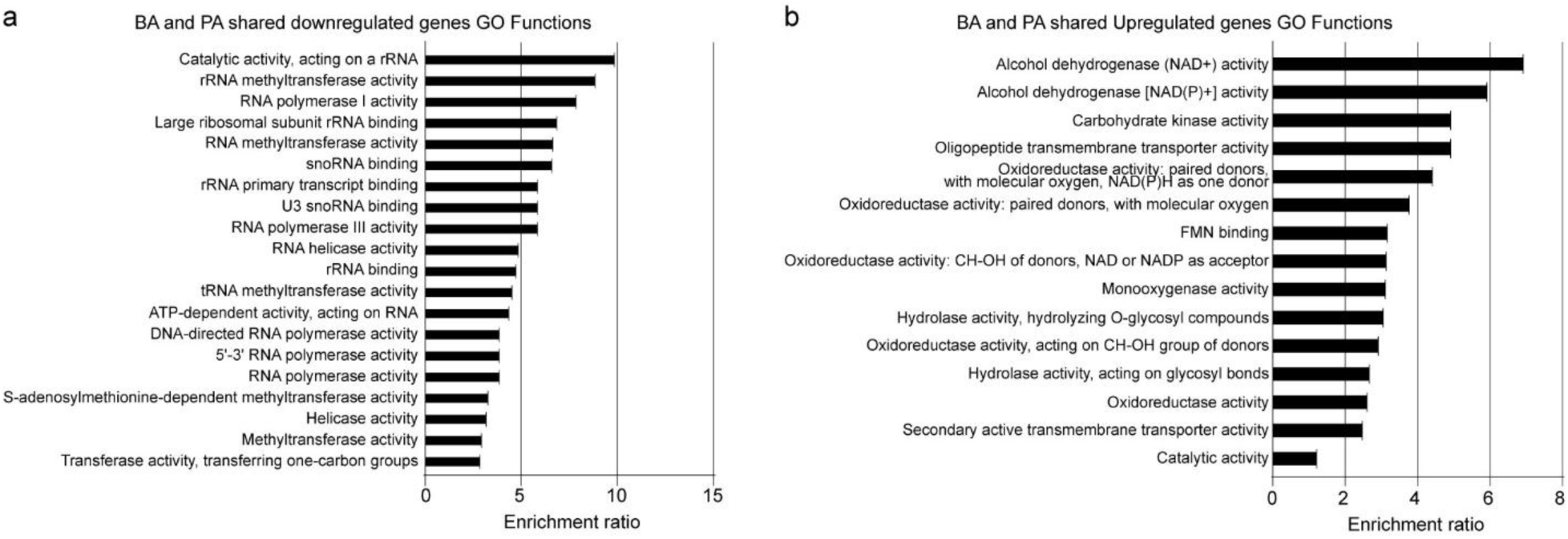
GO Functions of the differentially expressed genes shared by BA and PA treated *C. albicans*. **(a)** Pathway enrichment analysis (Gene Ontology, top 20 biological functions; Bonferroni-adjusted *p* < 0.05) of BA/PA-shared downregulated genes. **(b)** Pathway enrichment analysis (Gene Ontology, top 20 biological functions; Bonferroni-adjusted *p* < 0.05) of BA/PA-shared upregulated genes.

**Supplemental Figure 3.**
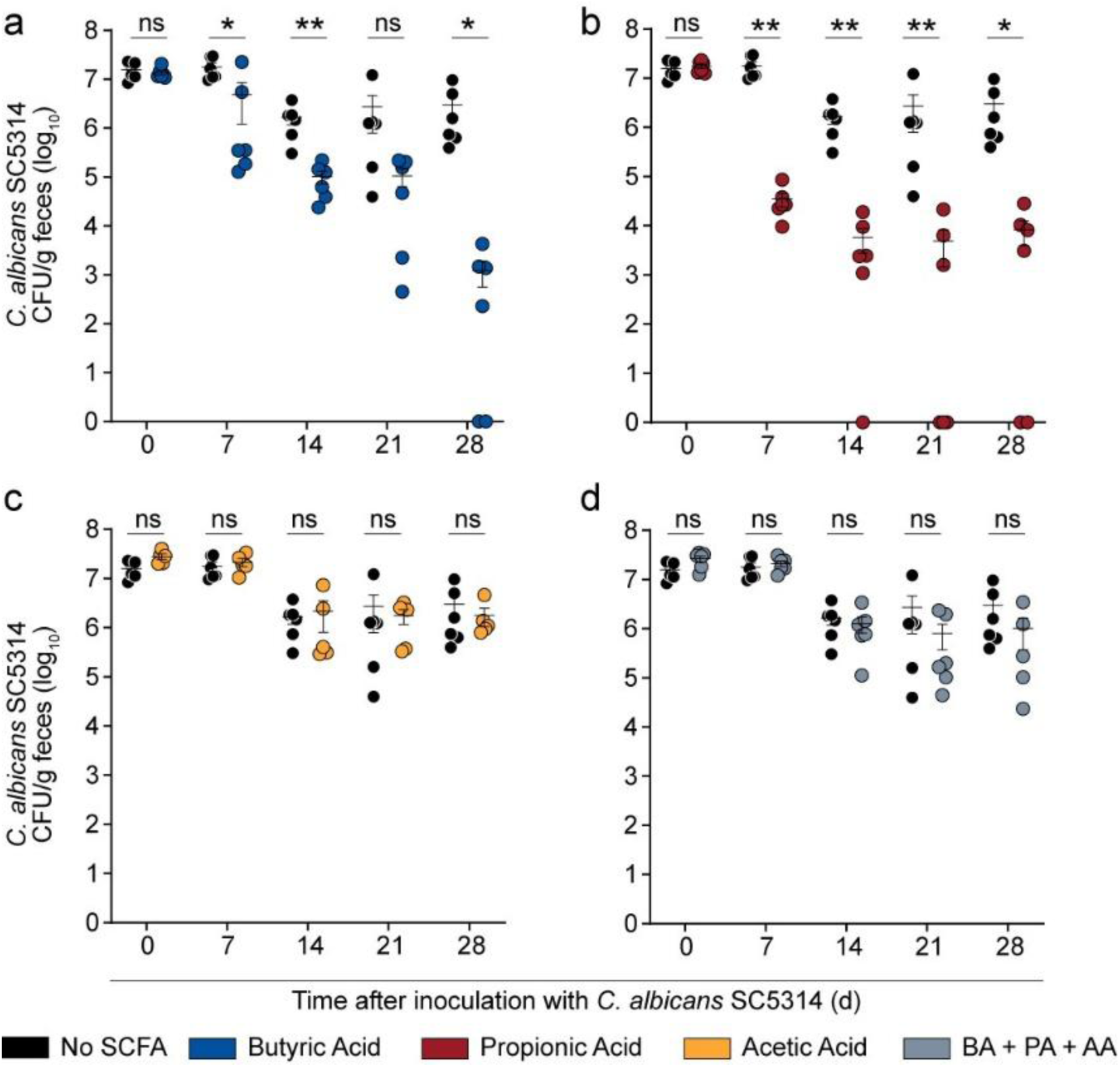
SCFA reduce *C. albicans* colonization in C3H/HeN mice. C3H/HeN mice (female, 6-8 weeks, Envigo) received penicillin/streptomycin in drinking water for 7 days to deplete commensals, were orally gavaged with 2 × 10^8^ CFU *C. albicans* SC5314, and then switched to sterile, pH-adjusted (5.0) drinking water ±150 mM SCFA ad libitum. *Ca* GI colonization levels, as determined by cultured enumeration of fecal homogenates on YPD with gentamicin and vancomycin. **(a)** Butyric acid **(b)** Propionic acid **(c)** Acetic acid **(d)** 42 mM butyric + 24 mM propionic and 84 mM acetic acid (150 mM total) Data from two independent experiments; n = 6 mice per group. Points = results from individual animals. Bars = mean ± SEM. Mann-Whitney test. \**P* < 0.05, \*\**P* < 0.01, ns, not significant.

**Supplement Figure 4.**
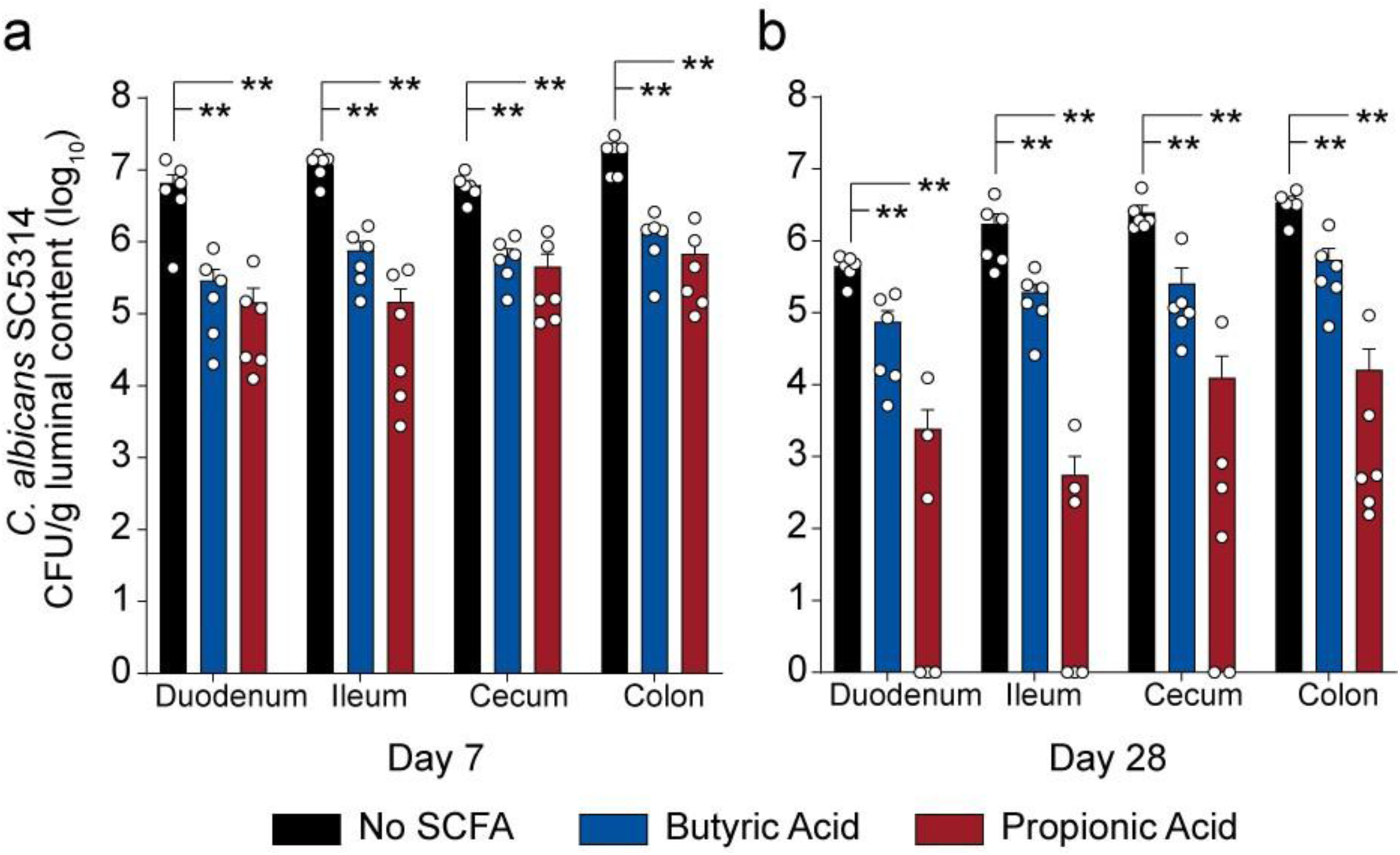
SCFA reduces *C. albicans* colonization in different GI segments. **(a-b)** Mice C57/BL6J (female, 6-8 weeks, Jackson) received penicillin/streptomycin in drinking water for 7 days to deplete commensals, were orally gavaged with 2 × 10^8^ CFU *C. albicans* SC5314, and then switched to sterile, pH-adjusted (5.0) drinking water ±150 mM SCFA ad libitum. *Ca* luminal content colonization levels, as determined by cultured enumeration of luminal content homogenates on YPD with gentamicin and vancomycin. Content collected from the duodenum, ileum, cecum and distal colon. **(a)** day 7 **(b)** day 28 Points = results from individual animals. Bars = mean ± SEM. Two independent experiments were performed with final n=6 per group. Mann-Whitney test. \*\**P* < 0.01.

**Supplemental Figure 5.**
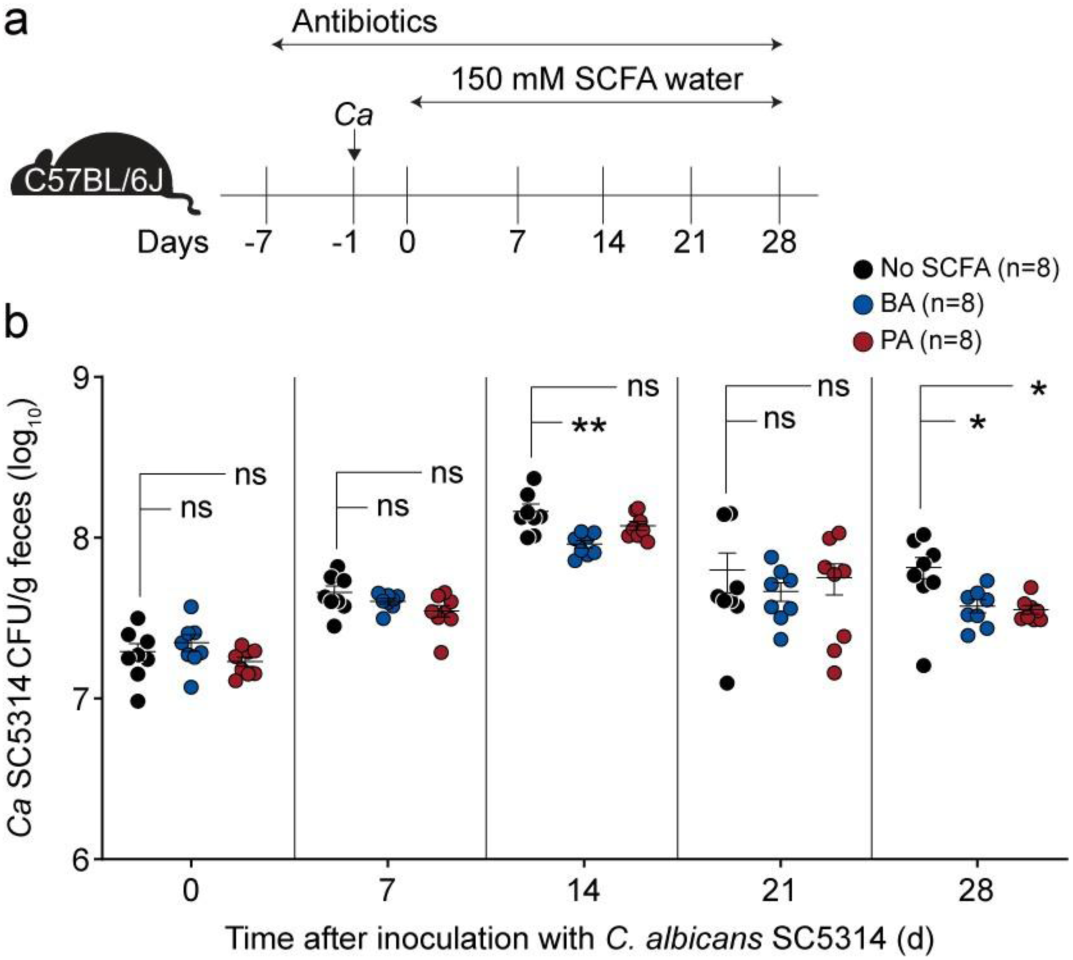
*C. albicans* colonization in mice receiving concomitant SCFA and antibiotic treatment. **(a)** Experimental schema for assessing continuous antibiotics and SCFA treatment in conventional C57/BL6J mice (female, 6-8 weeks, Jackson). Mice received penicillin/streptomycin in drinking water for 7 days to deplete commensals, were orally gavaged with 2 × 10^8^ CFU *C. albicans* SC5314, and then switched to sterile, pH-adjusted (5.0) drinking water ±150 mM SCFA ad libitum. Colonization was monitored for 28 days. **(b)** *Ca* GI colonization levels, as determined by cultured enumeration of fecal homogenates on YPD with gentamicin and vancomycin. Data from two independent experiments; n = 8 mice per group. Points = results from individual animals. Bars = mean ± SEM. Mann-Whitney tests. \**P* < 0.05, \*\**P* < 0.01, ns, not significant.

**Supplemental Figure 6.**
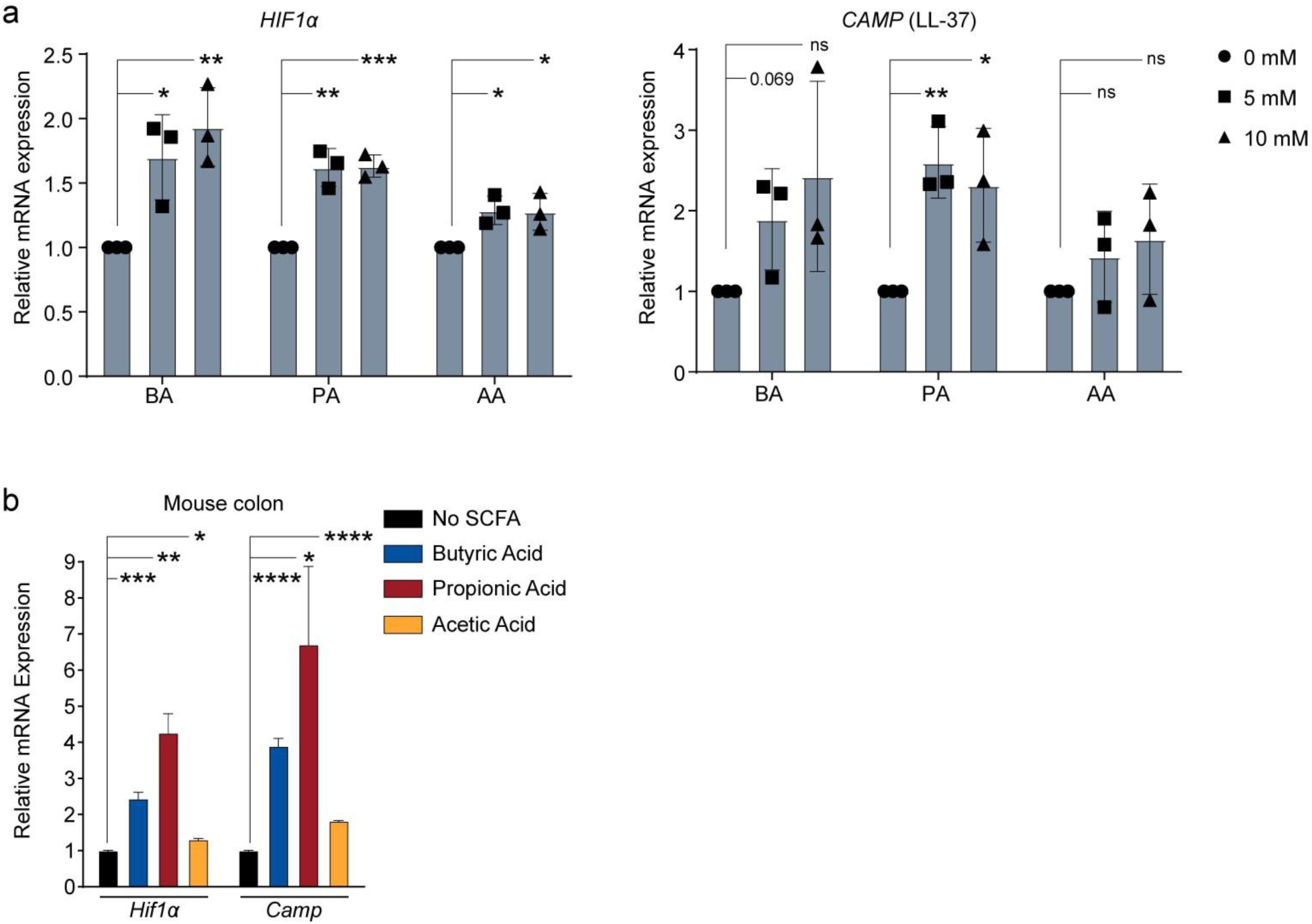
SCFAs induce HIF-1α and CRAMP gene expression in human and mouse colonocytes. **(a)** HIF-1α (*HIF1α*) and LL-37 (*CAMP*) gene expression in human colonocytes (HT-29) with SCFA treatment, in pH-adjusted media. Experiment was performed in triplicate with three technical replicates per condition. Unpaired *t*-test. \**P* < 0.05, \*\**P*< 0.01, *** P < 0.001, ns, not significant. **(b)** Gene expression of *Hif1α* and *Camp* from the colons of C3H/HeN (female, 6-8 weeks, Envigo) mice that received ±50 mM SCFA ad libitum. Data from two independent experiments, n=4. Unpaired *t*-test. \**P* < 0.05, \*\**P* < 0.01, *** P < 0.001, **** P < 0.0001, ns, not significant

**Supplemental Figure 7.**
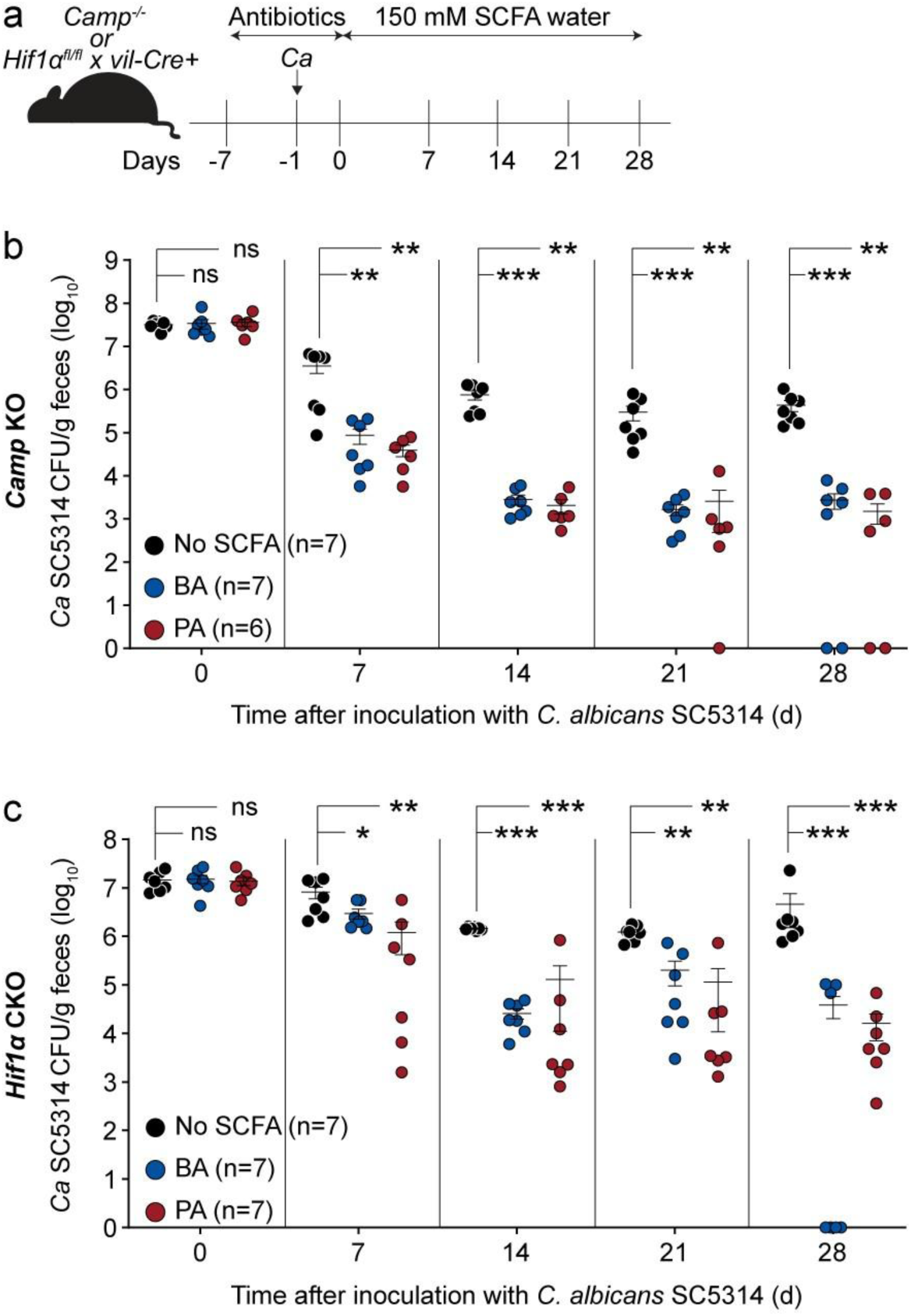
SCFA reduces *C. albicans* GI colonization in *Camp* KO and *Hif1α* CKO mice. **(a)** Experimental schema for assessing SCFA treatment in conventional *Camp* knockout (KO) (female, 6-8 weeks old, Jackson) or *Hif1α* conditional knockout (CKO) mice (female, 6-8 wks old, Jackson). Mice received penicillin/streptomycin in drinking water for 7 days to deplete commensals, were orally gavaged with 2×10^8^ CFU *C. albicans* SC5314, and then switched to sterile, pH-adjusted (5.0) drinking water ±150 mM SCFA ad libitum. Colonization was monitored for 28 days. **(b)** *Ca* GI colonization levels in *Camp* KO mice, as determined by cultured enumeration of fecal homogenates on YPD with gentamicin and vancomycin. Data from two independent experiments; n = 6–7 mice per group. **(c)** *Ca* GI colonization levels in *Hif1α* CKO mice, as determined by cultured enumeration of fecal homogenates on YPD with gentamicin and vancomycin. Data from two independent experiments; n = 6–7 mice per group. Points = results from individual animals. Bars = mean ± SEM. Mann-Whitney tests. \**P* < 0.05, \*\**P* < 0.01, \*\*\**P* < 0.001, ns, not significant.

**Supplemental Figure 8.**
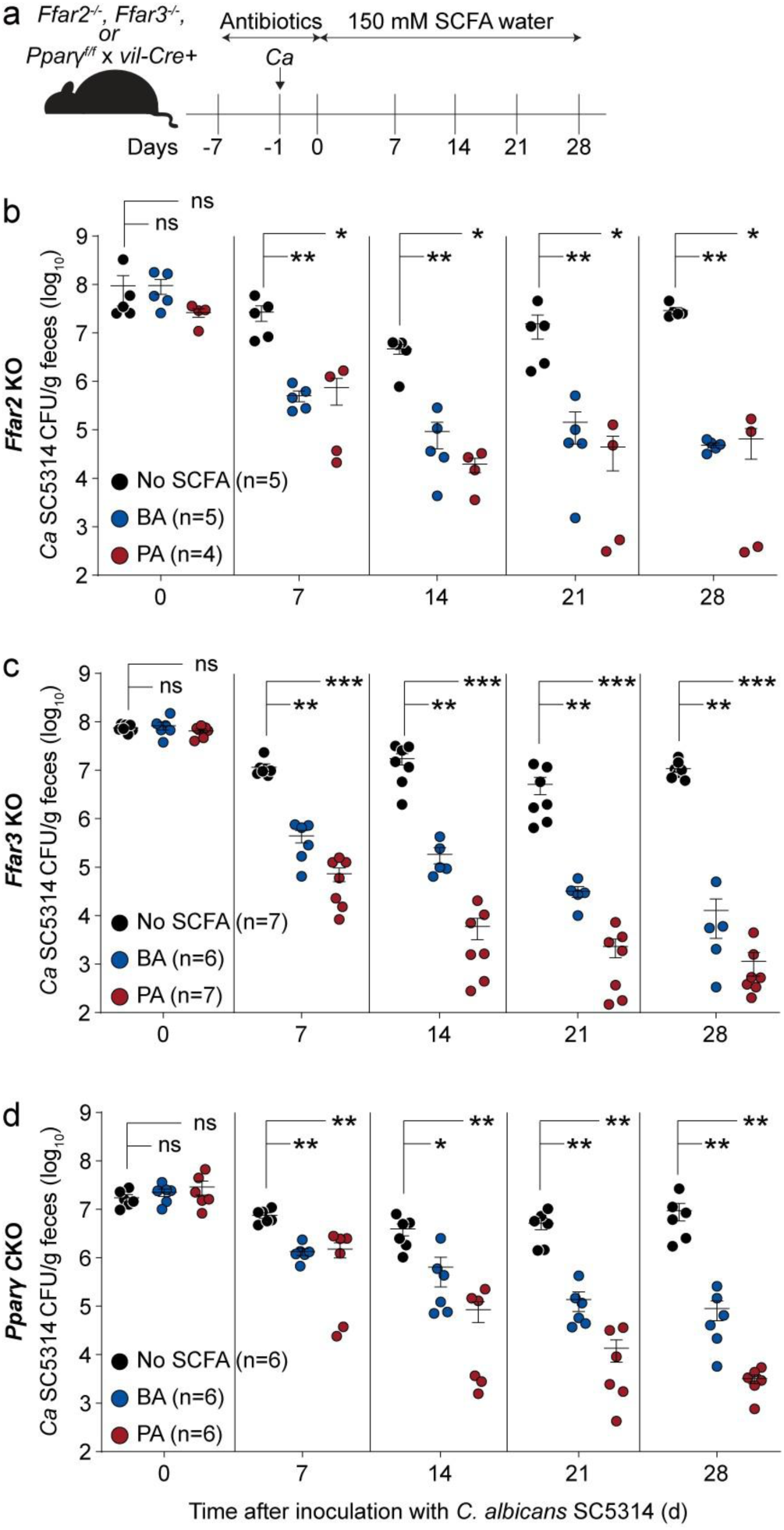
SCFA reduces *C. albicans* GI colonization in absence of host SCFA receptors or sensors. **(a)** Experimental schema for assessing SCFA treatment in conventional *Ffar2* KO (female, 6-8 weeks old, MMRRC)*, Ffar3* KO (female, 6-8 wks old, RIKEN) and *PPARγ* (female, 6-8 weeks old, Jackson) CKO mice. Mice received penicillin/streptomycin in drinking water for 7 days to deplete commensals, were orally gavaged with 2×10^8^ CFU *C. albicans* SC5314, and then switched to sterile, pH-adjusted (5.0) drinking water ±150 mM SCFA ad libitum. Colonization was monitored for 28 days. **(b)** *Ca* GI colonization levels in *Ffar2* KO mice, as determined by cultured enumeration of fecal homogenates on YPD with gentamicin and vancomycin. Data from two independent experiments; n = 4-5 mice per group. **(c)** *Ca* GI colonization levels in *Ffar3* KO mice, as determined by cultured enumeration of fecal homogenates on YPD with gentamicin and vancomycin. Data from two independent experiments; n = 6–7 mice per group. **(d)** *Ca* GI colonization levels in *PPARγ* CKO mice, as determined by cultured enumeration of fecal homogenates on YPD with gentamicin and vancomycin. Data from two independent experiments; n = 6 mice per group. Points = results from individual animals. Bars = mean ± SEM. Mann-Whitney tests. \**P* < 0.05, \*\**P* < 0.01, \*\*\**P* < 0.001, ns, not significant.

